# Contact networks of small mammals highlight potential transmission foci of *Mammarenavirus lassaense*

**DOI:** 10.1101/2025.02.25.639449

**Authors:** David Simons, Ravi Goyal, Umaru Bangura, Rory Gibb, Ben Rushton, Dianah Sondufu, Joyce Lamin, James Koninga, Momoh Foday, Mike Dawson, Joseph Lahai, Rashid Ansumana, Elisabeth Fichet-Calvet, Richard Kock, Deborah Watson-Jones, Kate E. Jones

## Abstract

Lassa fever (*Mammarenavirus lassaense*; LASV), is an endemic zoonosis in West Africa. Human infections arise from rodent-to-human transmission, mainly from the synanthropic reservoir *Mastomys natalensis*. In Sierra Leone, small-mammal communities vary across land use gradients, shaping LASV transmission risk. How anthropogenic environments facilitate the rodent-rodent interactions remains poorly understood.

We sampled small mammals over 43,266 trap nights in Sierra Leone’s LASV-endemic Eastern Province, detecting 684 rodents and shrews. To assess potential transmission, we constructed space-sharing networks from co-trapping events within species-specific home range radii. These networks approximated shared space use, allowing comparison of encounter patterns across habitats.

Network topology varied significantly by land use. Village networks were the most connected (highest average degree), whereas agricultural communities supported the most species (higher rarefied richness) and were the most fragmented (higher modularity). Notably, the probability of intraspecific space sharing among *M. natalensis* was highest in agriculture, suggesting land use modulates key intra-specific transmission pathways.

LASV seroprevalence was 5.7% community-wide, with antibodies in nine species. We found no statistically significant association between overall seroprevalence and land use or aggregate network structure (mean degree). However, predictive modeling for *M. natalensis* indicated that higher individual degree is associated with seropositivity, suggesting complex, scale-dependent relationships. These findings show that simple ecological drivers do not fully explain LASV exposure, highlighting the importance of species-specific behaviors (i.e., *M. natalensis* clustering in agriculture) and the multi-host serological landscape in assessing transmission risk.

## Introduction

Lassa fever caused by *Mammarenavirus lassaense* (LASV) is a rodent-associated zoonosis endemic to West Africa, with an estimated 100,000-900,000 infections annually^1,2^. While outbreaks in Nigeria are routinely reported, cases in other endemic countries - Guinea, Liberia and Sierra Leone - are sporadically documented^3–5^. In Sierra Leone disease outbreaks frequently go undetected, consistent with findings of up to 80% LASV seropositivity among human communities in regions previously not considered endemic^6^. Human infections typically result from transmission from rodent hosts, with limited subsequent human-to-human transmission^7^. Understanding LASV transmission in endemic settings requires a detailed characterization of small-mammal community interactions, through which pathogen transmission occurs and is maintained.

The primary reservoir of LASV, *Mastomys natalensis* is a native synanthropic rodent species, widespread across sub-Saharan Africa. Pathogen challenge studies on *M. natalensis* colonies suggest that acute infection does not significantly alter rodent behavior or cause clinical pathology^8–10^. LASV is transmitted through both direct contact (e.g., superficial wounds caused by infected conspecifics) and indirect contact (e.g., exposure to contaminated environments), at low infectious doses^9^. Vertical transmission (mother-to-pup) is also thought to be an important mechanism for viral persistence and spread^10,11^. Infected adult rodents exhibit detectable viral RNA as early as 3 days post-infection, with viral loads peaking within 1-2 weeks and resolving by 40 days^9^. Among individuals infected within the first 2 weeks of life viral RNA is detectable up to 16 months post infection^10^. The transient nature of acute infection - outside of neonatal infections - has led many studies to focus on LASV-specific antibody detection, rather than markers of acute infection, such as viral RNA^12–14^.

The antibody response dynamics to LASV in rodents are not yet well understood. Based on data from a similar arenavirus, Morogoro virus, seroconversion is expected to occur by 7 days post-infection, with antibodies (e.g., IgG) persisting beyond the decline of detectable RNA^11^. LASV-infected rodents are presumed to develop lifelong immunity to reinfection upon seroconversion, however, the efficacy of neutralizing antibodies is unclear and the role of immune or partially immune individuals in transmission networks is not known^9,10,15^. Although antibody-based studies have limitations, the higher prevalence of LASV seropositivity compared to acute infections provides valuable insights into viral dynamics within endemic regions. For example, a recent study in Bo district, Sierra Leone, reported a 2.8% LASV IgG seroprevalence among rodents and shrews, compared to a 0.3% prevalence of acute infection detected via PCR, underscoring the challenges of identifying acute infections in small-mammal populations^16^.

While *M. natalensis* is considered the primary reservoir species of LASV, evidence of LASV infection has been found in 15 other small-mammal species in endemic regions: five identified with acute infection and ten showing previous infection based on serological evidence.^14,17–19^. The contribution of these additional species to pathogen transmission into human populations - and their role in viral transmission or maintenance within their species communities - remains unclear. In species-rich environments, both direct and indirect contact among small mammals may result in incidental infections of non-reservoir species, which are subsequently detected through surveillance. Incidental infections of non-primary reservoir species may have little impact on viral transmission or maintenance^20^. Alternatively, these species could facilitate the transfer of LASV across landscapes, linking geographically isolated *M. natalensis* populations and causing reintroduction of the virus into reservoir species populations^21,22^. To account for this uncertainty we refer to *M. natalensis* as the primary reservoir species and other small-mammal species previously found to be infected as hosts. Increasing recognition of multi-species host systems in zoonoses underscores the importance of expanding surveillance efforts to the wider community in which the host resides to better understand pathogen prevalence and dynamics^23,24^.

Understanding the structure of small-mammal contact networks in LASV-endemic regions may offer valuable insight into the structural drivers of transmission, even in the absence of direct observations of infection. While prior studies have described rodent diversity and abundance, these do not capture interaction patterns relevant for pathogen spread^16,25,26^. Spatial proximity can be used as a proxy for potential contact and, when analyzed as networks, can help identify structural features, such as connectivity, inter- and intra-specific mixing, and network fragmentation that influence pathogen transmission. In contexts where direct behavioral observation or tagging of individuals is infeasible, such as post-Ebola Sierra Leone, proximity-based approaches provide a pragmatic alternative to infer likely transmission pathways^27^.

Small-mammal communities in LASV-endemic regions are structured along anthropogenic land use gradients^14,28^. As such, the risk of Lassa fever outbreaks in human populations is expected to correlate with these gradients^6,29,30^. Within human-dominated land use types, the prevalence of typically synanthropic rodent hosts of LASV is anticipated to be higher due to increased food availability, shelter, and reduced predation pressure^31–33^. These factors influence rodent abundance and population dynamics which in turn may promote greater pathogen persistence, as observed in several other rodent-associated zoonosis systems^34–37^. Understanding how small-mammal contact networks, alongside small-mammal occurrence and abundance, vary along these anthropogenic gradients could reveal potentially distinct pathogen transmission networks in different habitats. We hypothesize that both contact frequency and overall network connectivity increase in human-dominated habitats where resources are concentrated, leading to enhanced potential for pathogen transmission.

In this study, we define “contact” as the inferred co-occurrence within a spatial and temporal window (i.e., proximate capture locations within four trap-nights), consistent with prior spatial proximity models^38,39^. While this does not imply physical contact or indirect contact suitable for transmission, it serves as a proxy for shared habitat use and potential indirect or environmental contact relevant to LASV transmission. Interaction networks derived from wildlife data have previously been employed to study pathogen transmission. These networks are particularly valuable for investigating the role of community structure and the impact of contact rate heterogeneity between species in multi-host pathogen systems^40–42^.

We leverage rodent and shrew trapping data collected over three years in the Lassa fever-endemic region of the Eastern Province, Sierra Leone. We characterize potential interactions - both direct and indirect - within these communities as a network, where the nodes represent rodents or shrews, and the connections (or edges) between them represent potential interactions. We further hypothesize that the spatial clustering of conspecifics and the increased abundance of commensal species in anthropogenically dominated environments will lead to higher intra-specific contact rates compared to inter-specific contact rates within these communities. We use network analysis to explore how contact patterns vary along an anthropogenic land use gradient, focusing on *M. natalensis* while also assessing the roles of other potentially important, highly connected species (“hubs”). Finally, we report the prevalence of antibodies against LASV among individual small mammals in the study region and investigate the association between contact rates with seropositivity.

## Methods

### Study area

Rodent trapping surveys were conducted between October 2020-April 2023 within and around four village study sites (Baiama, Lalehun, Lambayama, and Seilama) in the Lassa fever endemic zone of the Eastern Province of Sierra Leone (Figure 1A). Surveys were conducted within trapping grids along a land use gradient of anthropogenic disturbance comprising forest, agriculture (including fallow and currently in-use areas), and villages (within and outside of permanent structures) (see Supplementary Information for study site and trap grid locations). One study site (Lambayama) was more developed than the other villages due to its proximity to the province capital (Kenema), we refer to this site as peri-urban with the others termed rural. Trapping survey sessions occurred four times annually with two trapping surveys in each of the rainy and dry seasons (May to November and December to April, respectively), producing up to a total of 10 trapping sessions in each village over the study period.

**Figure 1.**
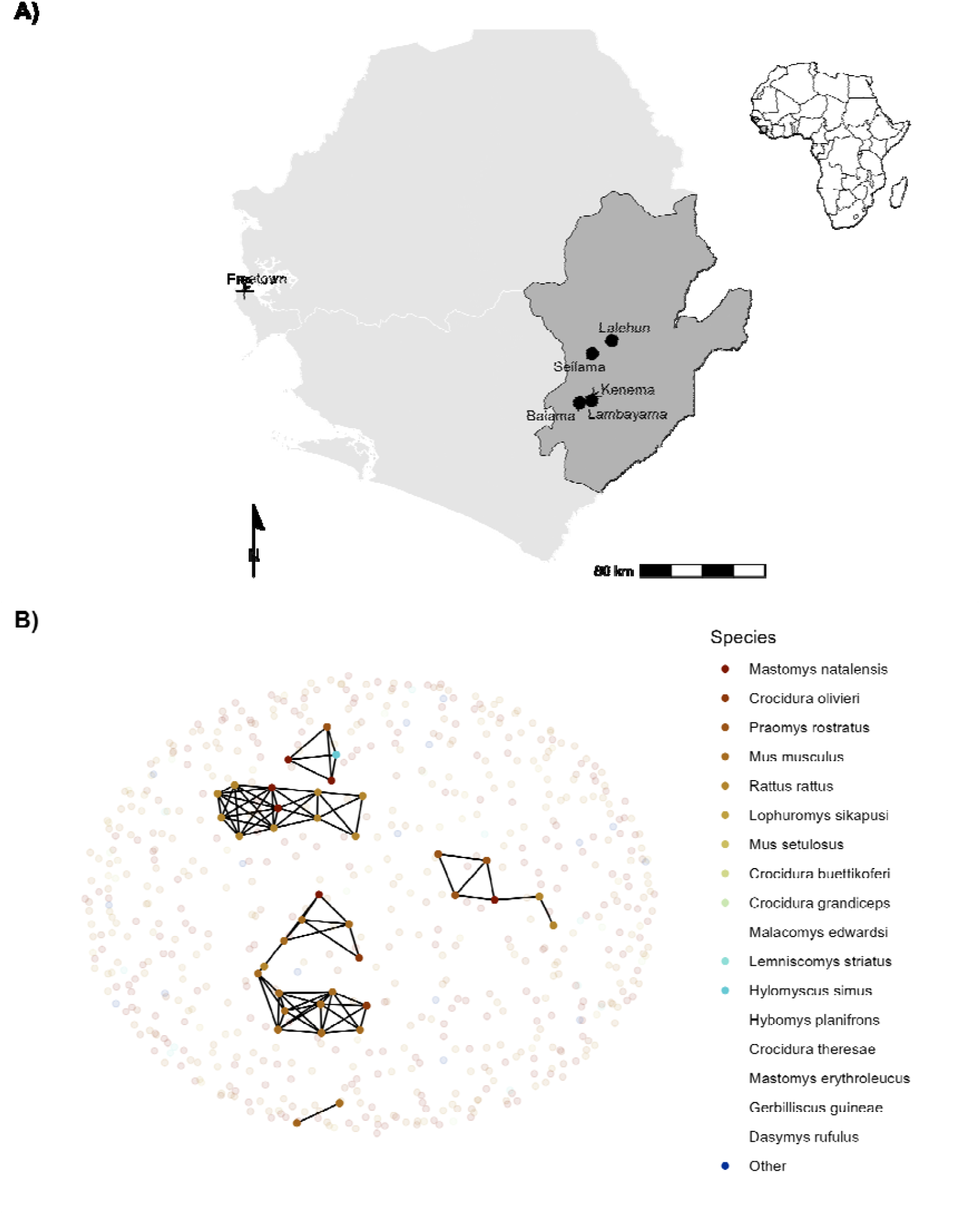
A) Location of village study sites (circles) in the Eastern Province of Sierra Leone. Kenema, the largest city in the province, and the national capital, Freetown, are also shown (+). The inset map highlights the location of Sierra Leone within West Africa. B) An example of a rodent contact network derived from trapping data during visit 5 in village land use. Each colored node represents an individual small mammal, with lines (edges) indicating inferred contacts between individuals. The number of edges connected to a node represents its degree. Betweenness reflects the importance of a node in connecting different parts of the network. Pale nodes indicate unobserved individuals, for whom contacts (edges) were not recorded. Species listed in the legend without colors were never detected in the shown land use type (village).

Alt text: [A two-panel figure. Panel A is a map of Sierra Leone, marking the four village study sites in the Eastern Province, with an inset showing Sierra Leone’s location in West Africa. Panel B is a network diagram showing potential contacts between individual small mammals, where nodes colored by species represent animals and lines represent shared space use.]

Study sites were selected to represent the range of land use in the Eastern Province of Sierra Leone, considering both accessibility throughout the year and acceptability of the study protocol to the village communities (Supplementary Information). At each trapping grid 49 Sherman traps (7.62cm x 8.89cm x 22.86cm) (H.B. Sherman Traps, Tallahassee, USA), were arranged in a 7 x 7 grid, with traps placed 7 meters apart in a grid conforming to the local landscape (median trapping grid area = 1,672 m^2^). In permanent structures, trap placement deviated from the grid structure. At each visit, permanent structures were randomly selected from a grid projected over the village area, with four traps placed within each structure. The location of each trap within the grid was geolocated. Traps were baited with a locally produced mixture of oats, palm oil and dried fish. Each morning, traps were checked, closed for the day, and re-baited in the evening. Each trapping survey session consisted of four consecutive trap-nights (TN) at each trapping grid within the village study site. Rodents and shrews were associated with the coordinates of the trap they were detected. Geospatial processing was performed using the sf package in R (version 4.1.2)^43,44^.

All small mammals were handled by trained researchers using appropriate personal protective equipment. Animals were sedated using halothane and euthanized according to established protocols^45^. Morphological measurements and samples of blood and tissue (skin, liver and spleen) were collected. The study was approved by the Clinical Research Ethical Review Board and Animal Welfare Ethical Review Board of the Royal Veterinary College, UK (URN: 2019 1949-3), and the ethics committee of Njala University, Sierra Leone. The study adhered to national and institutional ethical guidelines for the humane treatment of animals and safe handling of zoonotic pathogens. Trapping and sampling protocols were designed to minimize stress to the animals and reduce the risk of pathogen exposure to researchers and communities. Community engagement sessions were conducted to ensure understanding and acceptance of the study objectives, and all work complied with the principles outlined in the ARRIVE guidelines (v2)^46^. All carcasses were incinerated to mitigate pathogen transmission risks.

### Species identification

Species identification was performed in the field based on external morphological characteristics, including body length, tail length, ear length, and pelage coloration, following the taxonomic keys of Happold and Kingdon and Monadjem *et al*.^47,48^. Field identification was supplemented by molecular methods to confirm species identity for individuals identified as *Mastomys sp*., *Mus sp*., *Rattus sp*. and *Crocidura sp*. alongside a random subset of remaining individuals (50% of remaining samples).

All samples remained in Sierra Leone and were stored at −20°C until processing. Genomic DNA was extracted using QIAGEN DNeasy kits as per the manufacturers instructions^49^ (Supplementary Information). DNA extracts were amplified using platinum *Taq* polymerase (Invitrogen) and *cytochrome B* primers^16^. DNA amplification was assessed through gel electrophoresis, and successful amplification products were Sanger sequenced (performed by Eurofins Genomics). The sequences were attributed to rodent species using the BLAST program, comparing the obtained sequences to *cytochrome B* records in the NCBI database (accessed 2023-06-30)^50^.

### Serological Analysis

Serological status of trapped rodents and shrews was determined using the BLACKBOX® LASV IgG ELISA Kit (Diagnostics Development Laboratory, Bernhard Nocht Institute for Tropical Medicine), which has been validated for rodent samples^51,52^. The protocol is reproduced in Supplementary Information. Briefly, 1µL of whole blood was inactivated and mixed with the provided sample dilution buffer (1:50). Where whole blood was unavailable (21 samples, 3%), blood was eluted from dried blood spots stored on filter paper by incubating with phosphate-buffered saline containing 0.08% Sodium Azide and 0.05% Tween-20^53^. Samples, alongside negative and positive controls, were incubated on the ELISA kit plates for 24 hours at 4-8 °C in a wet chamber. Following incubation, the plates were washed and incubated for a further hour with 1:10,000 diluted HRP-labelled streptavidin. A final wash was performed prior to the addition of 100µL of 3,3’,5,5’-Tetramethylbenzidine (TMB) substrate to wells, with incubation for 10 minutes. The colorimetric reaction was stopped by adding 100µL of a stop solution.

A deviation from the kit protocol occurred due to local ELISA plate reader limitations. We measured the optical density (OD) at 450nm and 630nm, as opposed to 450nm and 620nm but this was not expected to have an important effect on absorbance patterns, as advised by the manufacturer. The index value was calculated by subtracting OD_630_ from OD_450_ and dividing by the cut-off value (the mean values of the negative controls + 0.150). Samples were classified as positive if the index value was greater than or equal to 1.1, negative if the index value was less than or equal to 0.9, and inconclusive if the index value was between 0.9 and 1.1. Inconclusive results were retested.

The prevalence of seropositive individuals is reported aggregated by species. A Bayesian logistic regression model was constructed, using the brms package, to estimate the Odds Ratio (OR) of seropositivity for each species compared to *M. natalensis*, which served as the reference species^54^. Specifically, a Bernoulli regression with normal, uninformative priors for population-level effects was used, with a binary dependent variable for seropositivity and an independent variable of small mammal species. Only species with more than 10 individuals assayed for antibodies to LASV were included in this model. Posterior distributions are presented in graphical format, alongside the posterior mean and 95% Credible Interval (CrI). Unlike frequentist approaches, Bayesian inference does not rely on *p*-values; rather, statistical support for an effect is assessed based on the posterior distribution, with associations interpreted in terms of the central tendency (e.g., OR) and whether the CrI excludes the null value (OR = 1).

In addition to the species-level seroprevalence analysis, we conducted *post-hoc* exploratory analyses to assess differences in LASV seroprevalence by village and land use type. These analyses employed Bayesian logistic regression models analogous to those used in species-level comparisons, with results interpreted cautiously given their non-pre-specified nature.

### Community Diversity Analysis

To compare small-mammal diversity across land use types, it was necessary to account for the different number of animals captured in each habitat. We therefore used individual-based rarefaction (IBR), a statistical technique that standardizes species richness estimates as if an equal number of individuals had been sampled in each habitat. This approach allows for a robust comparison of alpha diversity by controlling for unequal sampling effort.

Using the mobr package in R, we calculated the expected species richness for each habitat rarefied to a common sample size of 10 individuals, which was the minimum number of captures in any single habitat^55,56^. To further investigate the ecological processes shaping these communities, we compared these rarefied richness values to the expectations of a null model. This comparison allowed us to assess whether the observed species abundance patterns deviated from random community assembly, for instance, to detect potential dominance effects where one or a few species reduce overall community evenness and richness.

### Contact Network Construction

Species space-sharing networks were reconstructed from the trapping data. Capture-mark-recapture (CMR) methods have previously been used to identify space-sharing by individuals^39,57,58^. In our study system, a CMR design was not feasible due to study communities concerns around the risk of releasing an infected animal. Therefore, we considered that spatial and temporal co-occurrence proxies direct or indirect contacts with other small mammals^38^. We assumed these potential contacts were sufficient to transmit LASV if they were trapped within a species specific buffer zone centered on the location of the trap during the same 4 trap night session. A key assumption underlying this approach is that an individual was trapped at the center of their home range^58^.

We obtained species specific buffer radii for the primary analysis using a hierarchical approach that prioritized the most robust data available for each species. Our primary source was systematically compiled data within the HomeRange R package (version 1.0.2), from which we extracted median radii at the species level where possible, including for *M. natalensis* (median home range radii = 10.6m), *Mus musculus* (9.6m), *Lemnisomys striatus* (8m), *Lophuromys sikapusi* (8.4m), and *Rattus rattus* (29.3m)^59^. For detected species with no species level home range data, *Praomys rostratus* (7.2m), *Mus setulosus* (9.6m), *Crocidura spp*. (7.2m) and *Hylomyscus simus* (7.7m) we used the genus level median home range. Where no data were available we obtained home range radii from single-study estimates in the literature. This provided values for *Malacyoms edwardsi* (35m), *Gerbilliscus guineae* and *Hybomys planifrons* (both 22m)^60–62^. For *Dasymys rufulus*, which lacked any of the above data, we assigned the median of all previously determined species-level radii (11 m). To assess the impact of these assumptions, we performed pre-specified sensitivity analyses by applying uniform buffer radii of 15, 30, and 50 meters to all species.

Networks were constructed from observed animals (nodes) and the presence or absence of contact between them (edges). Data were aggregated by land use type and sampling visit, producing a potential 32 distinct networks from 201 trapping grid, village and visit combinations (see Figure 1B for an example network). However, as no rodents were detected in three networks derived from forest sites, only 29 networks were used in subsequent analysis.

### Network Structure and Mixing Pattern Analysis

We analyzed the structure of the space-sharing networks at three distinct levels: 1) the overall network within each land use type, 2) the individual animals (nodes) within those networks, and 3) species-level roles, including connectivity patterns (degree distribution, hub identification) and mixing dynamics (inter-vs. intra-specific contacts).

First, to characterize the overall networks, we calculated global metrics for each land use type (village, agriculture, and forest). These included the total number of nodes (animals) and edges (inferred contacts), as well as the network-wide mean degree and betweenness centrality. We then used a Kruskal-Wallis test to formally compare the distributions of these metrics across the three land use types. For metrics with a significant overall difference, we performed a Dunn’s *post-hoc* test with Bonferroni correction to identify which specific pairs of habitats differed.

Second, to evaluate the roles of individual animals, we calculated two key node-level centrality measures: degree centrality, the number of connections for each animal, as a direct measure of its contact frequency, and betweenness centrality, to identify individuals acting as critical bridges within the network. We then investigated species-specific interaction patterns by analyzing the degree distribution for each species to identify key hub species (i.e., those with notably high connectivity) in each habitat (Figure 3). To formally test for differences in connectivity across land uses, we used a Wilcoxon rank-sum test to compare the network-level mean degree for key abundant species.

To quantify mixing patterns, we also calculated the proportion of each species’ contacts that were intraspecific (with its own species) versus interspecific (with other species), providing a measure of network homophily across the land use gradient.

### Modeling Contact Probability in *Mastomys natalensis*

To examine the association between land use and species with the probability of a contact between two individuals, we modeled these contacts using Exponential-Family Random Graph Models (ERGMs)^63^. The analysis was limited to *M. natalensis*, the primary rodent host of LASV. Estimation of ERGM parameters provide an OR for the probability of an edge in a network – conditional on the rest of the network - based on network properties included in the model and nodal attributes. Within our trapping grids, only a subset of all individuals are detected in traps. Including unobserved individuals - and thus, unobserved contacts - enhances the interpretability and generalizability of the network models. This approach allows for a more accurate estimation of the total population size by accounting for missing data, thereby making the network models more representative of the entire population from which the analytic sample was derived.

Previous analysis of our study system suggests that the probability of detecting a rodent at each trap is less than 10% for 4 trap nights, provided that the species is present in the trapping grid^28^. To estimate the abundance of individuals of each species within a trapping grid, we modeled abundance (i.e., total population size) from repeated count data using an N-mixture model implemented in the unmarked R package (version 1.2.5)^64,65^. The latent abundance distribution was modeled using Poisson, negative binomial or zero-inflated Poisson random variables. The abundance model included the number of trap nights and season as replicate-dependent detection covariates, as well as location (rural vs. peri-urban setting) and land use type (forest, agriculture or village) as occurrence covariates.

To select the most appropriate model for each species, we compared the Akaike Information Criterion (AIC) of the Poisson, negative binomial, and zero-inflated Poisson models. The best-fitting model was then used to derive the estimated abundance. The median estimated abundance from the distribution produced for each trapping grid was used to estimate the number of unobserved individuals in each network, aggregated by land use type (Supplementary Figures 2.1-2.12). The number of observed individuals was subtracted from the predicted abundance to derive the number of unobserved individuals for each species. These unobserved individuals were explicitly set to have missing (i.e., unobserved) edge values.

**Figure 2.**
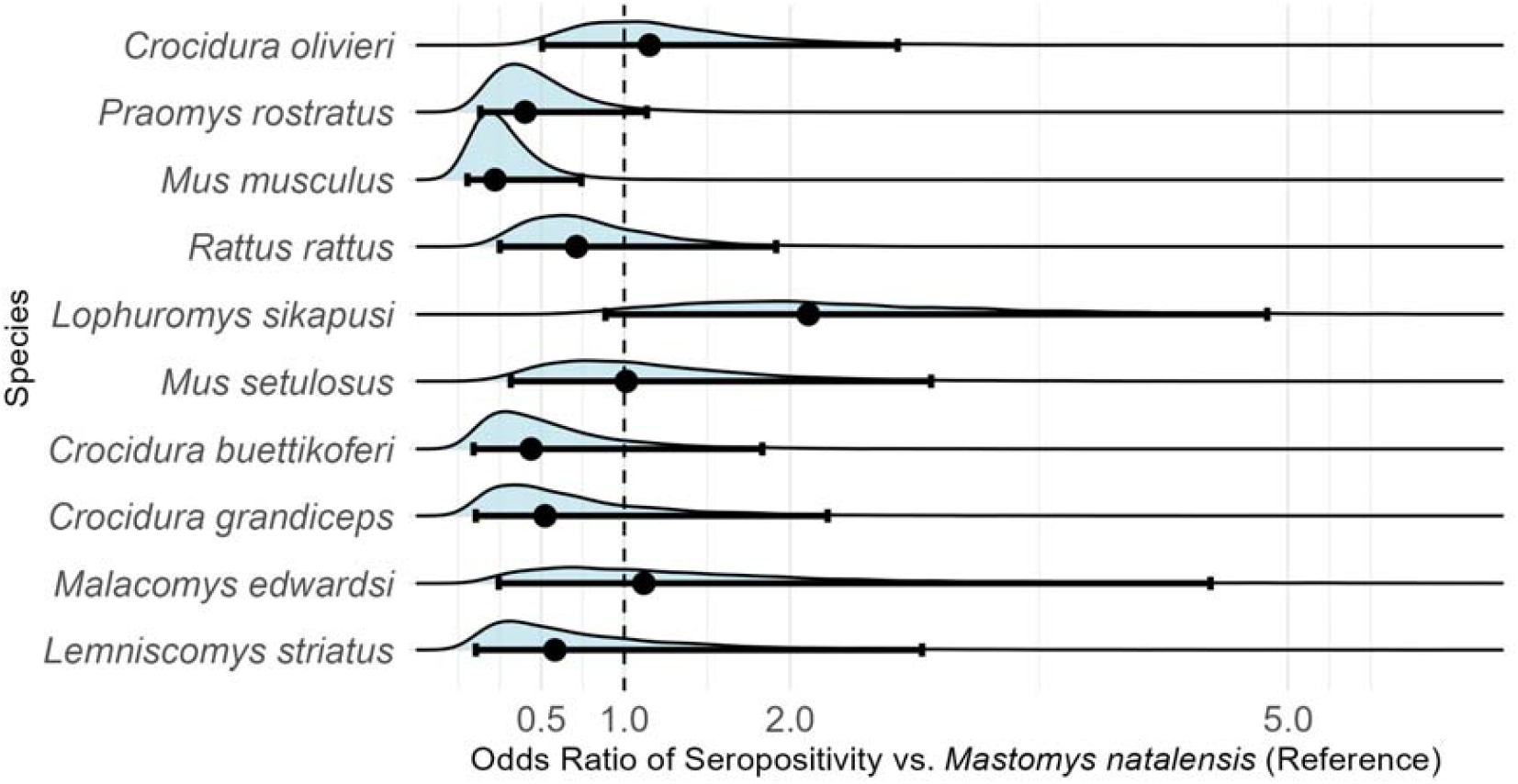
Odds Ratios of seropositivity to LASV among small-mammal species, compared to Mastomys natalensis. Only species with more than 10 individuals assayed for antibodies to LASV were included in this analysis

**Figure 3.**
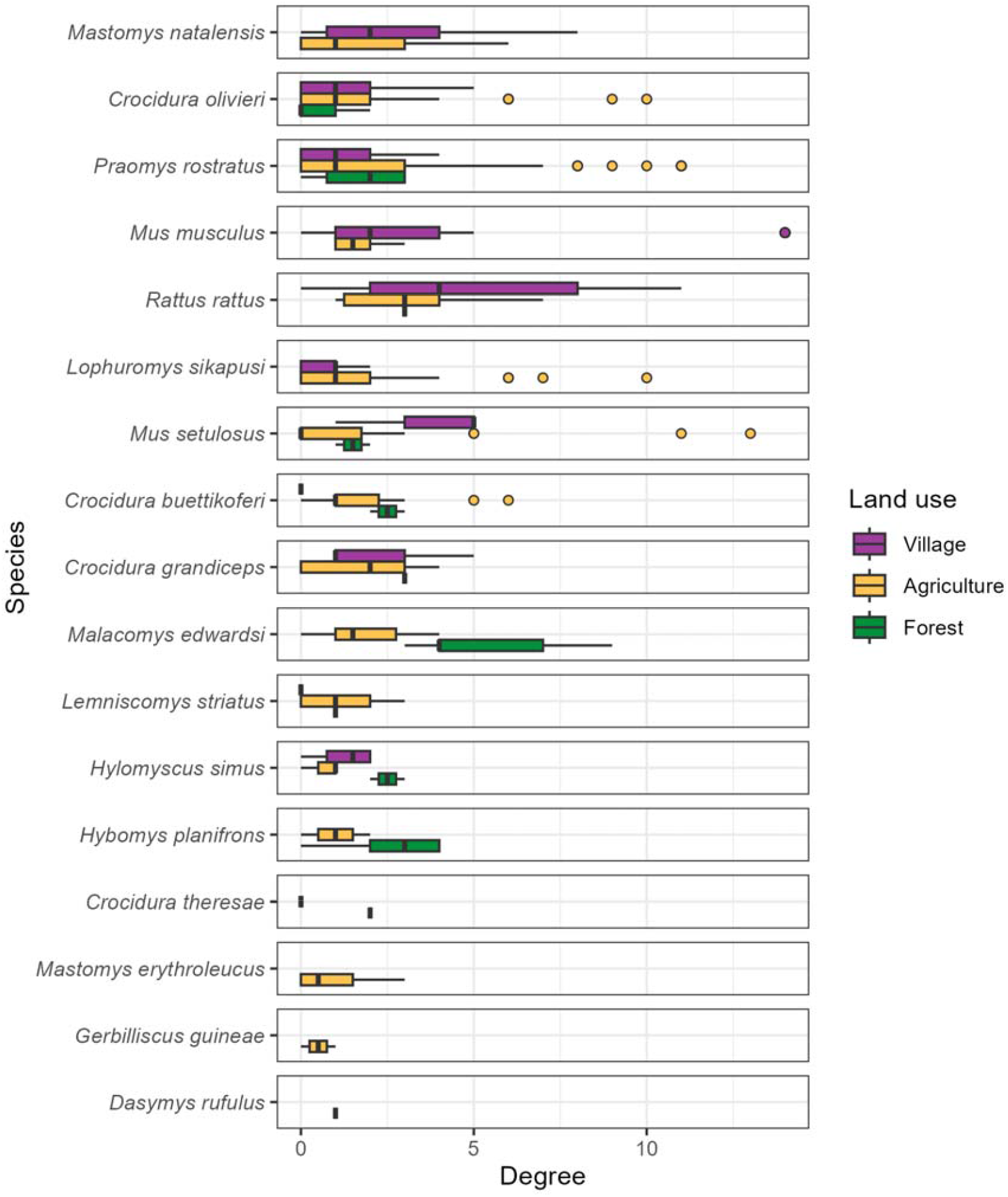
The degree (number of contacts) of individual small mammals stratified by species and land use type. Boxes contain the median and inter-quartile range of the degree distribution. Whiskers include the upper and lower quartile with outliers shown as points.

Finally, the constructed adjacency matrices were converted to networks using the network R package (version 1.13.0.1) for subsequent ERGM modelling^66^ (Figure 1B) and Supplementary Figures 3.1-3.3).

ERGMs were specified for each of our inferred contact networks to compare the probabilities of edges forming based on rodent characteristics (i.e., species). The general model is shown in Equation 1:

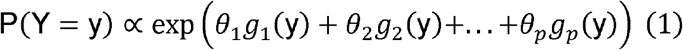

Where *p* is the number of terms in the model, and the values of the coefficients *θ* represent the size and direction of the effects of the covariates *g*(y) on the overall probability of an edge being present in the network. At the edge level the expression for the probability of the entire graph can be re-expressed as the conditional log-odds of a single edge between two nodes (a contact between two rodents) as shown in Equation 2.

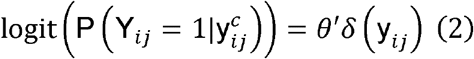

Where Y_*ij*_ is the random variable for the state of the node pair *ij* and 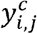 signifies all dyads in the network other than *y*_*i,j*_. *θ*′ is the coefficient for the change production of an edge between the two nodes conditional on all other dyads remaining the same 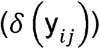.

ERGMs are implemented using the ergm package (version 4.3.2) in R^67^. Three terms were included in the final ERGM to model the probability of the formation of ties (Equation 3). The first term (edges), describes the density of the network, representing the probability of a tie being observed in the network. The second term (species) represents the conditional probability of a tie forming, conditional on the species of the nodes. The third term (species homophily) accounts for intraspecific tie formation among rodent individuals (i.e., the conditional probability of two individuals of the same species forming a tie).

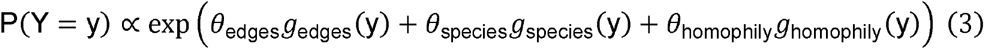

ERGMs were implemented on the individual networks for each land use type at each visit. The effect sizes from each model were pooled through random-effects meta-analysis, stratified by land use, to produce a land use-specific summary effect size for each coefficient^68^. Inclusion in meta-analysis was restricted to ERGMs that produced stable estimates for each of the model terms (i.e., sufficient detections of *M. natalensis* within the network). Random-effects models were conducted using the `metafor` package (version 4.0.0) in R^69^. Heterogeneity across the models was assessed using the *Q*-test and the restricted maximum-likelihood estimator (*τ*^2^) with a prediction interval for the true outcomes produced^68,70^. The *Q*-test assesses whether there is greater variability among effect sizes than expected by chance, with a significant result indicating substantial heterogeneity. The *τ*^2^ statistic estimates the between-study variance, quantifying the degree of heterogeneity rather than just testing for its presence. Weights for each network included in meta-analysis were assigned using inverse-variance weights^71^.

The presence of influential networks was assessed using Cook’s distance, for models including influential networks leave-one-out sensitivity analysis were performed^72^. Forest plots were generated to visualize the summary OR of the probability of a tie for each model term, stratified by land use type.

Models with unstable estimates for the species homophily term were not included in the random-effects meta-analysis. No contact networks from forest land use contributed to meta-analysis as no *M. natalensis* were detected in these settings. Five models from agricultural settings and six from village settings were included in meta-analysis.

### Analysis of Network Centrality and Serostatus

To investigate pathogen transmission within our networks, using seropositivity as a proxy for prior exposure to LASV, we first report the small-mammal species found to contain individuals that were seropositive for LASV. We then compared the nodal degree of seropositive and seronegative individuals using a Wilcoxon rank-sum test with continuity correction^73^. This analysis was repeated stratified by species to assess whether contact rates were associated with an individual being seropositive. Finally, we compared the node-level betweenness of seropositive and seronegative individuals to determine whether an individual’s position within a structured contact network was associated with prior exposure to LASV.

### Modelling Seropositivity Risk Factors in *M. natalensis*

We next tested whether an individual *M. natalensis*’s risk of being LASV-seropositive was influenced by its position in the contact network. Specifically, we investigated the effects of two key properties: its total number of contacts (degree) and its tendency to connect with conspecifics (homophily). Because our trapping data provides only one realization of a dynamic social system, we used a simulation-based approach to account for this structural uncertainty. Based on the network formation rules derived from the ERGMs described previously, we generated 50 plausible network simulations for each trapping grid that contained at least one seropositive *M. natalensis*.

Using these simulated networks, we fitted a Bayesian generalized linear mixed-effects model (GLMM) to assess the association between network position and serostatus. The GLMM predicted LASV serostatus as a function of degree, homophily, and their interaction, with random intercepts for each original trapping grid and simulation replicate to account for the nested data structure. The model was specified using the following formula:

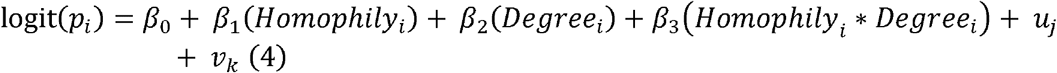

Where *u*_*j*_ (the random effect for each network) is drawn from a Normal distribution with a mean of 0 and a variance 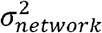, and *v*_*k*_ (the random effect for each simulation) is drawn from a Normal distribution with a mean of 0 and a variance 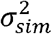. For each *M. natalensis* individual in each simulation, we calculated its degree and a node-level homophily score (defined as the proportion of its direct contacts that were with other *M. natalensis*). Evidence for an association was assessed using the posterior distributions of the model coefficients.

## Results

A total of 684 small mammals were captured over 43,266 trap-nights, representing 17 species (13 rodent and 4 shrew species). *M. natalensis* was the most commonly detected species (N = 113, 16.5%), followed by *Crocidura olivieri* (N = 105, 15.3%) and *Praomys rostratus* (N = 102, 15%) (Table 1). Rarefied species richness (*S*), standardized to 10 individuals, was 5.86 in agricultural habitats, 5.13 in forests, and 4.36 in village settings. Observed species richness was 9 in agricultural sites, 6 in villages and 3 in forests. In village habitats, rarefied richness values were consistently lower than the expectations of null models of species abundance distributions.

**Table 1:** The number of individuals detected and antibodies to LASV among those individuals.

| Species | Order | Individuals (N) | LASV Antibody detected (%) | Percentage of all positive individuals | OR (95% CrI) |
| --- | --- | --- | --- | --- | --- |
| *Mastomys natalensis* | Rodentia | 113 | 11 (9.7%) | 28.2% | ref |
| *Crocidura olivieri* | Eulipotyphla | 105 | 8 (7.6%) | 20.5% | 1.15 (0.51-2.65) |
| *Praomys rostratus* | Rodentia | 102 | 2 (2%) | 5.1% | 0.4 (0.13-1.14) |
| *Mus musculus* | Rodentia | 90 | 0 (0%) | 0% | 0.22 (0.05-0.74) |
| *Rattus rattus* | Rodentia | 88 | 4 (4.5%) | 10.3% | 0.71 (0.25-1.92) |
| *Lophuromys sikapusi* | Rodentia | 57 | 8 (14%) | 20.5% | 2.11 (0.89-4.88) |
| *Mus setulosus* | Rodentia | 43 | 3 (7%) | 7.7% | 1.01 (0.32-2.85) |
| *Crocidura buettikoferi* | Eulipotyphla | 23 | 0 (0%) | 0% | 0.44 (0.09-1.83) |
| *Crocidura grandiceps* | Eulipotyphla | 15 | 0 (0%) | 0% | 0.52 (0.11-2.23) |
| *Malacomys edwardsi* | Rodentia | 11 | 1 (9.1%) | 2.6% | 1.12 (0.24-4.53) |
| *Lemniscomys striatus* | Rodentia | 11 | 0 (0%) | 0% | 0.58 (0.11-2.8) |
| *Hylomyscus simus* | Rodentia | 9 | 0 (0%) | 0% |  |
| *Hybomys planifrons* | Rodentia | 7 | 1 (14.3%) | 2.6% |  |
| *Mastomys erythroleucus* | Rodentia | 4 | 1 (25%) | 2.6% |  |
| *Crocidura theresae* | Eulipotyphla | 3 | 0 (0%) | 0% |  |
| *Gerbilliscus guineae* | Rodentia | 2 | 0 (0%) | 0% |  |
| *Dasymys rufulus* | Rodentia | 1 | 0 (0%) | 0% |  |

### LASV Seroprevalence Across Species and Habitats

Antibodies to LASV were identified in 39 rodents and shrews (39/684, 5.7%) from 9 species, including *M. natalensis* (11/39, 28%), *C. olivieri* (8/39, 21%), *Lophuromys sikapusi* (8/39, 21%) and *Rattus rattus* (4/39, 10%) (Table 1).

In a Bayesian model with *M. natalensis* as the reference, the OR for LASV seropositivity was highest in *L. sikapusi* (OR = 2.11, 95% Credible Interval (CrI) = 0.89-4.88). However, like most other species, its credible interval included 1.0, indicating substantial uncertainty. In contrast, *M. musculus* had the lowest odds of being seropositive (OR = 0.22, 95% CrI = 0.06–0.74), and its CrI did not include 1.0, suggesting stronger evidence for reduced odds compared to *M. natalensis*.

Alt text:[A forest plot comparing the odds ratios of LASV seropositivity for several small mammal species against the reference species, *Mastomys natalensis*. Each species has a point estimate and a 95% credible interval. *Lophuromys sikapusi* shows a higher odds of being seropositive (OR 2.09), while *Mus musculus* shows lower odds (OR 0.22). The credible intervals for most species are wide and cross the null value of 1, indicating uncertainty.]

Antibody positive small mammals were detected in three of the study villages, Lalehun (N = 18, 46%), Seilama (N = 12, 31%) and Baiama (N = 9, 23%). No positive individuals were detected in Lambayama, the most urbanized village study site. Antibody positive individuals were detected during all study visits except visit 9 (2023-February). The proportion of antibody positive among all captures was 6.3% (24/379) in agricultural sites, 5% (13/261) in villages, and 4.5% (2/44) in forests.

In exploratory *post-hoc* models, the OR for seropositivity was 2.57 in Lalehun (95% CrI = 1.28-5.15), 1.55 in Baiama (95% CrI = 0.68-3.44), and 0.21 in Lambayama (95% CrI = 0.05-0.67), relative to Seilama as the reference village. The OR for agricultural land use was 1.27 (95% CrI = 0.68-2.42), and 0.85 (95% CrI = 0.23-2.59) in forests compared to villages. As the 95% CrIs for several comparisons (specifically, Baiama village, and agricultural and forest land uses) included 1.0, we did not find strong evidence for a difference in seroprevalence in these settings compared to their respective references.

### Structure of Small-Mammal Contact Networks

The number of individuals (nodes) was highest in agricultural settings (n = 379) compared to villages (n = 261) and forests (n = 44). While mean network degree was numerically highest in village settings (mean = 3.39), followed by forest (mean = 2.23) and agricultural settings (mean = 1.84), these differences were not statistically significant (Kruskal-Wallis, *H* = 3.05, *p* = 0.22). Similarly, we found no significant difference in mean betweenness centrality across land use types (*H* = 2.64, *p* = 0.27).

In contrast, network modularity differed significantly among habitats (*H* = 15.9, *p* < 0.001). *Post-hoc* tests revealed that agricultural networks were significantly more modular than both village networks (*p* = 0.02) and forest networks (*p* < 0.001). The difference in modularity between village and forest networks was not statistically significant. This indicates that small-mammal communities in agricultural settings form more fragmented, clustered interaction networks.

At the species level, degree centrality varied by habitat, with distinct hub species emerging in different settings (Figure 3). In villages, the synanthropic species *M. musculus* and *R. rattus* were highly connected, reaching maximum degrees of 14 and 11 with high mean degrees (4.1 and 4.7, respectively). In forests, the native species *Malacomys edwardsi* was a key hub, exhibiting the highest mean degree (5.4) and the maximum individual degree (9) for that habitat. By contrast, the principal reservoir *M. natalensis*, though abundant, showed more moderate connectivity with a lower maximum degree and lower mean degrees across habitats (mean = 2.3, max = 8 in villages and mean = 1.74, max = 6 in agriculture).

While the mean degree for several synanthropic species appeared numerically elevated in villages, formal comparisons of network-level mean degrees found no statistically significant differences in the degree distributions for species, including *M. natalensis* and *R. rattus*, across land use types (Wilcoxon rank-sum tests, *p* > 0.05 for all comparisons).

### Inter- and Intra-specific Mixing Patterns

Species with more detected individuals generally exhibited a greater number of inter-specific contacts (Pearson correlation, *r* = 0.73, *p* < 0.001, *n* = 15). For instance, the frequently detected species, *M. natalensis, P. rostratus* and *R. rattus* each had contacts with more than 11 other species. An exception to this trend was *M. musculus*, which, despite being the fourth most frequently detected species, only had observed contacts with four other species (Figure 4 and Supplementary Figures 4.1 and 4.2).

Intra-specific contacts were common for most species, but notable differences emerged across land use types. In agricultural settings, 46% of all contacts involving *M. natalensis* were intra-specific, even while interacting with 12 other species (Figure 4). In village settings, this proportion of intra-specific contact decreased to 28% (Supplementary Figure 4.2). Other species showed a higher degree of inter-specific mixing (e.g., *L. sikapusi* and *M. setulosus*).

**Figure 4.**
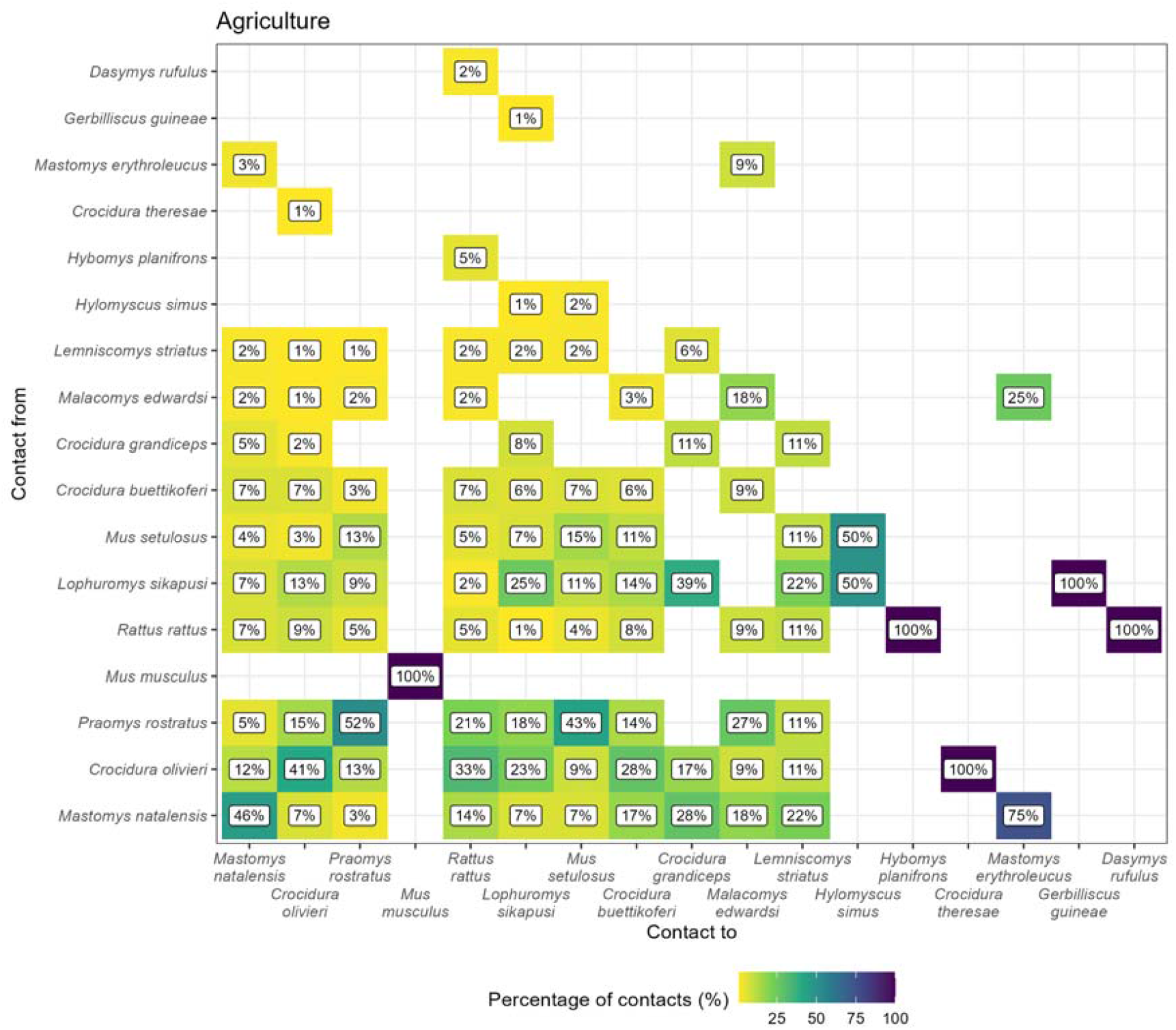
The proportion of contacts between individual small mammals in agricultural land use. Darker colors indicate increasing proportions of observed contacts to a species (Contact to) from named species (Contact from). Numbers in the cells correspond to the proportion of contacts to a species from a named species. For example, 45% of all contacts to Mastomys natalensis are from other M. natalensis while 9% of contacts are from Lophuromys sikapusi. Percentages sum to 100% in the `Contact to` axis, while they may exceed 100% in Contact from. Species are ordered by the total number detected in this study with M. natalensis (N = 113) in the bottom left.

### Modeling Contact Probability in *Mastomys natalensis*

Focusing on the primary reservoir, *M. natalensis*, we used a random-effects meta-analysis to pool the results from 11 individual ERGM models (6 village networks and 5 agricultural networks).

The overall probability of an inferred contact (the edges term) was low in both agricultural (OR = 0.04, 95% Confidence Interval = 0.03-0.07, *p* < 0.001) and village settings (OR = 0.15, 95% C.I. = 0.09-0.23, *p* < 0.001), with substantial heterogeneity observed between individual networks in both land use types (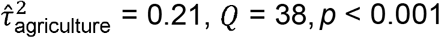 and 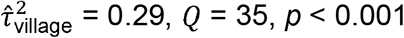) (Figure 5A).

**Figure 5.**
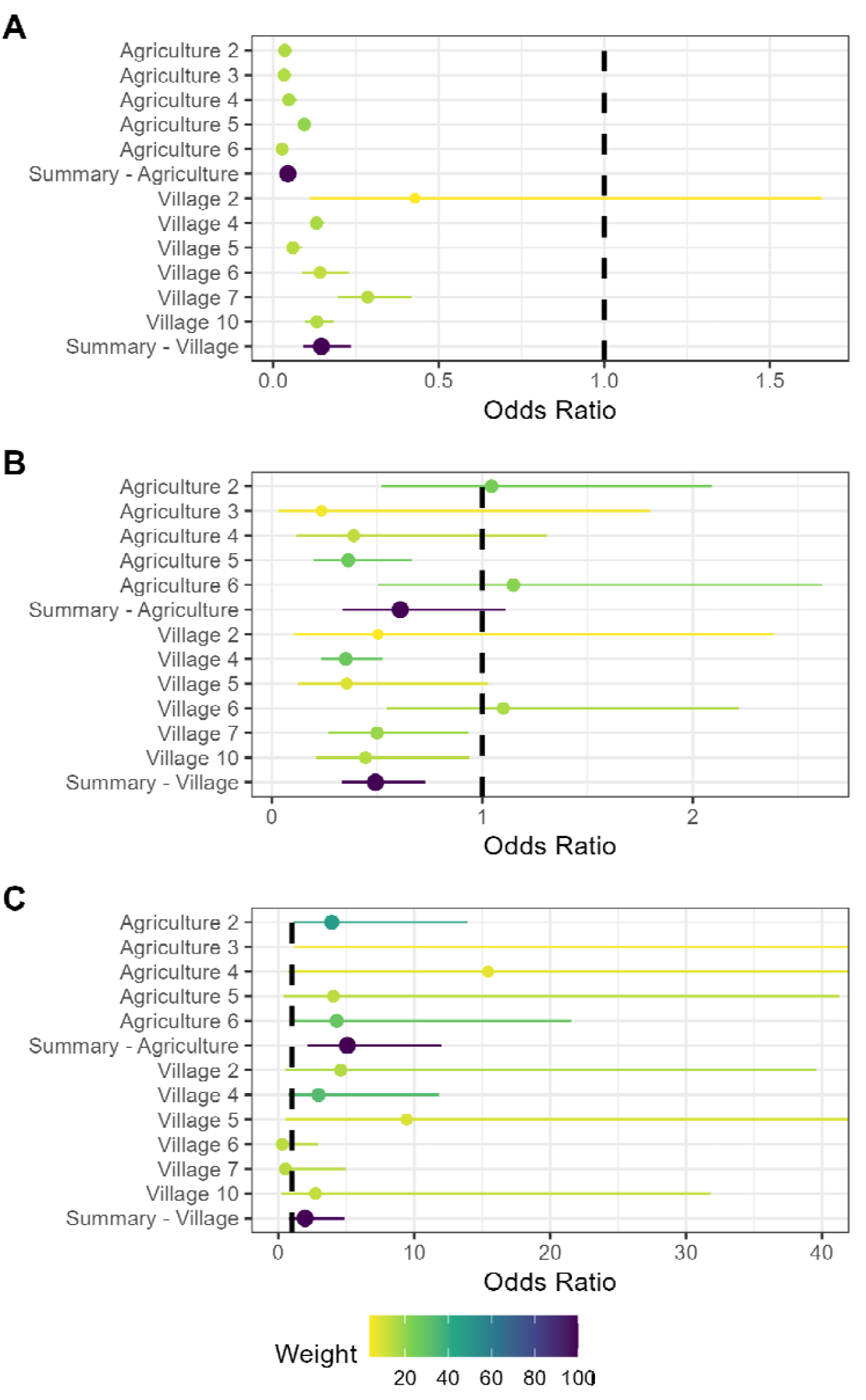
Random effects meta-analysis of ERGM network models reporting the odds of a contact being observed for M natalensis. A) The odds ratio of a contact being observed for M. natalensis in Agricultural or Village land use types. B) The odds ratio of a contact being observed between M. natalensis and an individual of a different rodent species. C) The odds ratio of a contact being observed between M. natalensis and another M. natalensis.

For inter-specific inferred contacts, *M. natalensis* had significantly reduced odds of forming a connection with another species in village settings (OR = 0.49, 95% C.I. = 0.33-0.73, *p* < 0.001). In agricultural settings the odds were also reduced, though this effect was not statistically significant (OR = 0.61, 95% C.I. = 0.33-1.11, *p* = 0.1) (Figure 5B). There was no substantial heterogeneity in inter-specific contact odds between networks (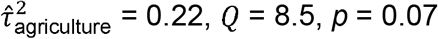 and 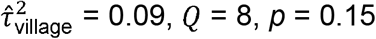).

Conversely, *M. natalensis* showed a strong and statistically significantly increase in the odds of intra-specific inferred contacts in agricultural settings (OR = 5.05, 95% C.I. = 2.14-12, *p* < 0.001), but not in village settings (OR = 1.96, 95% C.I. = 0.79-4.81, *p* = 0.15) (Figure 5C). Heterogeneity in intra-specific contact odds was low in both land use types (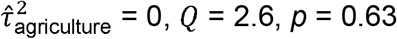 and 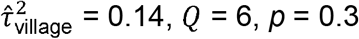).

Sensitivity analyses revealed no changes in the direction of effect sizes when altering the contact radius although the magnitude of the effect sizes varied (Supplementary Figures 5.1-3). Additionally, leave-one-out sensitivity analyses for influential networks did not indicate meaningful changes in the effect size magnitude or direction. These results support the robustness of the findings across space-sharing buffer area assumptions and community composition changes across visits.

### Observed Association Between Network Centrality and Serostatus

We first examined the observed association between an individual’s network centrality and its LASV serostatus. Overall, we found no significant difference in the mean degree of seropositive individuals (mean = 2, SD = 2.5) compared to seronegative individuals (mean = 2.5, SD = 3) (Wilcoxon rank-sum test, *W* = 11218, *p* = 0.29).

In species-specific analysis, this lack of a significant association held for most species, including the principal reservoir, *M. natalensis* (*p* = 0.43). However, for the invasive synanthrope *R. rattus*, seropositive individuals had a significantly higher mean degree than their seronegative counterparts (mean 7.75 vs. 4.24, *W* = 273.5, *p* = 0.034). This finding should be interpreted with caution, as it is based on a small number of seropositive individuals (n = 4). We found no significant difference in betweenness centrality between seropositive and seronegative individuals, either for the overall community or within any individual species (*p* > 0.05 for all tests).

### Modelling Predictors of Seropositivity Risks in *M. natalensis*

To move beyond correlation and to model the factors that predict seropositivity risk, we fitted a Bayesian generalized linear mixed-effects model for the primary reservoir, *M. natalensis*. This model, which included an interaction term, was favored over simpler models (Leave One Out comparison, ELPD difference >65).

In the final model, an individual’s odds of being LASV seropositive increased significantly with its total number of contacts (degree; OR = 2.36, 95% CrI: 1.98–2.81) and its proportion of conspecific contacts (homophily; OR = 10.22, 95% CrI: 5.39– 19.71). We also identified a strong negative interaction between degree and homophily (OR = 0.19, 95% CrI: 0.14–0.26).

## Discussion

In the Eastern Province of Sierra Leone, we found that the structure of small-mammal contact networks varied by habitat, though not as initially hypothesized. While we found no statistically significant difference in mean degree (connectivity) across land use types, key hub species differed distinctly. Invasive synanthropes like *R. rattus* and *M. musculus* were highly connected in villages, whereas the native species *M. edwardsi* emerged as a central hub in forests. The primary reservoir *M. natalensis* consistently showed a high probability of intra-specific contact in agricultural settings, suggesting these areas may host distinct transmission dynamics. Our most complex findings, however, relate to the link between network position and serostatus.

A key finding of our study is the divergence between our descriptive and modelling analyses of seropositivity. While our predictive model identified a higher degree as a significant risk factor for seropositivity in *M. natalensis*, our descriptive, non-parametric analysis found no significant overall difference in the mean degree of seropositive and seronegative animals. This divergence highlights the challenge of detecting clear disease signals in static, aggregated network data and suggests the underlying association is likely obscured by confounding factors that a simple correlational test cannot account for. The null result in our descriptive analysis is likely due to several mechanisms. Survival bias may play a key role, where highly-connected individuals face higher mortality or removal, thus masking the association between high degree and risk among the surviving population^74^. Most plausibly, the signal is obscured by the temporal dynamics of LASV. The virus likely persists through metapopulation dynamics and episodic local fadeout, and our serological data, aggregated over three years, smooths over these dynamics. In contrast, our model, which accounts for network structure, was able to detect the underlying signal that higher connectivity does indeed increase infection risk, a signal that is masked in a simple correlational analysis.

Our predictive model for *M. natalensis* also revealed a strong negative interaction between degree and homophily. This indicates that while having more contacts increased the odds of being seropositive overall, this effect was significantly weaker for individuals whose contacts were primarily with conspecifics. In other words, for highly connected animals, contacts with other species were more strongly associated with seropositivity than contacts with their own species. This finding could challenge the paradigm of *M. natalensis* as the sole driver of transmission, pointing towards a more complex, community-level maintenance system where inter-species spillover events are particularly important.

Analyzing the network structures highlights significant ecological heterogeneity across the land use gradient. The upper tail of the degree distribution was skewed towards individuals in villages and agriculture, underscoring the importance of individual-level variation and a few highly connected hub individuals not captured by aggregated metrics^75,76^. Furthermore, agricultural networks were more modular. While this suggests these communities may be fragmented into distinct clusters, a structure that could facilitate intense local transmission while slowing landscape-level spread, this finding should be interpreted with caution. The observed modularity may be partially influenced by the spatial arrangement of trapping grids in heterogeneous agricultural environments rather than reflecting social clustering alone. The high species richness and greater proportion of inter-specific space-sharing in agricultural areas also likely reflects edge effects that facilitate interactions between synanthropic and sylvatic species, creating opportunities for inter-species spillover^77,78^. Conversely, the high connectivity of invasive synanthropes in villages may contribute to the lower-than-expected species richness observed in those habitats through competitive exclusion or other mechanisms^79^.

The structure of *M. natalensis* interactions also varied by land use. In agricultural settings, *M. natalensis* exhibited significantly higher odds of intra-specific clustering, a pattern consistent with the species’ weak territoriality^80–82^. In villages, this strong tendency for intra-specific contact was not statistically significant. This dynamic could amplify intra-specific transmission chains in agricultural landscapes while potentially diluting transmission pathways in villages where inter-specific encounters may be proportionally higher. Movement between habitats by individuals driven by resource availability may further modulate these dynamics^83^.

The seroprevalence of LASV (5.7%), was consistent with prior estimates from Sierra Leone^16^. Our study included forest sites further from human habitation, yet the proportion of *M. natalensis* individuals testing positive was similar (∼9%). Our detection of LASV antibodies in nine distinct species, with *M. natalensis* comprising only 28% of seropositive individuals, confirms previous reports of LASV exposure across a range of small-mammal species in West Africa^12,18^. While seropositivity indicates past infection, it does not confirm reservoir competence or define a species’ role in active transmission. Nonetheless, this finding underscores the importance of considering a multi-host community perspective when studying LASV ecology^84^.

These complex ecological findings have direct public health implications. The distinct network structures suggest different human risk profiles by land use: the high intraspecific connectivity of *M. natalensis* in agricultural settings may amplify the virus within the primary reservoir, posing a direct spillover risk to the members of these communities that most utilize agricultural landscapes. In villages, the multi-host seropositivity and denser networks suggest a more diffuse risk to residents from a wider range of species. Furthermore, our finding that seropositive animals do not, on average, have higher connectivity implies that surveillance strategies focused only on the most socially central animals may be insufficient. A more comprehensive sampling approach is likely necessary to accurately assess community-level prevalence. Notably, we detected no seropositive animals in the most urbanized study site (Lambayama), suggesting factors associated with increased urbanicity, potentially including shifts in community composition towards invasive species or altered habitat structure, might disrupt local transmission cycles, though further investigation is needed. Ultimately, the confirmation of a multi-host serological landscape, despite low absolute numbers of seropositive individuals for many species, suggests that interventions targeting solely the primary reservoir may be insufficient or yield counterintuitive outcomes in complex ecological systems^85^.

Our study’s methodology rests on a series of assumptions that introduce cumulative uncertainty. Contacts were inferred from spatio-temporal co-occurrence, which assumes that proximity is a reliable proxy for interactions capable of transmission^38^. Our approach also assumes that a capture location represents a central point within an individual’s short-term activity space^58^. Furthermore, removal trapping may have created temporary spatial vacuums, and unobserved behavioral differences between captured and uncaptured animals could bias inferences^86,87^. While our study design (≥3 months between sessions), sensitivity analyses, use of species-specific home range radii, and leave-one-out meta-analysis sought to mitigate these issues, we acknowledge our results represent a model of a complex system. Ultimately, replicating the study across a greater number of sites would be valuable for evaluating the generalizability of these patterns.

In conclusion, this study highlights the variability in small-mammal contact networks across a land-use gradient in a Lassa fever-endemic region. While we found no direct association between land use and seroprevalence, our results reveal that the ecological drivers of transmission risk are complex and scale-dependent. The divergence between our descriptive and modeling results suggests that static snapshots may not fully capture dynamic disease processes. The serological evidence suggesting multi-host LASV exposure, alongside the habitat-specific roles of hub species underscore the importance of tailoring surveillance and control strategies to local ecological contexts to mitigate Lassa fever risks effectively.

