## Supplementary Information for "Contact networks of small mammals highlight potential transmission foci of *Mammarenavirus lassaense*"

### Rodent trapping and lab processing protocol

2025-09-26

**References** **13**

#### Preparation for field work

Before going to the field the following must be checked.

1. Adequate number of clean and functional Sherman traps are brought. You will need at least **324** to set up the correct number of grids.
2. Adequate sample pots and filter paper for rodent specimens.
3. Personal protective equipment for 16 sessions of sample processing.
4. Spare batteries for GPS devices.
5. Battery packs for electronic devices.
6. Portable freezer for rodent specimens, to be stored at Panguma Hospital Lab.
7. Paper copies of the data entry forms.

### Trapping protocol

This trapping protocol is based on previous rodent trapping studies conducted in Eastern Sierra Leone (Leski et al. 2015; Bangura et al. 2021). The approach has been modified to use trapping grids rather than lines.

#### Trap placement

Rodents are being trapped for an ongoing study of rodent population structures and assemblages in a Lassa fever endemic region of Eastern Sierra Leone. The study will occur over 2 years (November 2020 - February 2023). The primary outputs of the rodent trapping will be to identify to species level the small mammals in different land use types at different times of the year in addition to the presence of antibodies to Lassa fever virus. Trapping will occur in pre-specified study site locations every 4 months. Trapping is described at multiple spatial levels. The highest order is the study village, followed by a trapping grid and finally a trap location. Four study villages have been selected for ongoing work. The villages are Baiama, Lalehun, Lambayama and Seilama (Baiama; lat = 7.8375, long = -11.2683, Lalehun; lat = 8.1973, long = -11.0803, Lambayama; late = 7.8505, long = -11.1969, and Seilama; lat = 8.1224, long = -11.1936). At each village site up to 7 trapping grids have been designated. A trap grid contains 49 traps, in non-household settings each grid is a 7 x 7 regular grid of Sherman traps set up to 7 metres apart (Figure 1.). Within household settings four traps will be placed within each home. The land use of the grid and more specifically the trap location and its habitat structure will be noted at each study site visit. The coordinates of each trapping location are recorded to record the location at which each individual rodent was detected. At dusk all traps are baited and traps are primed, traps are checked each morning with traps containing rodents brought to the field processing site. The location of the trap is marked to allow replacement following processing of the rodent. All traps are closed during the day for re-baiting during the afternoon. Traps are set for four consecutive nights at each visit.

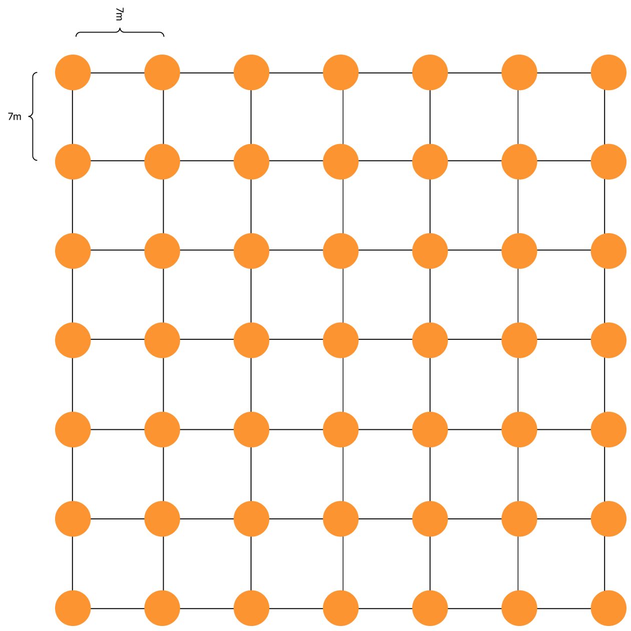

Figure 1: Structure of a 49 trap grid. Individual traps are placed 7 metres apart in a regular grid 7 by 7 grid. A solid orange circle represents the location of a single Sherman trap.

#### Rodent processing

Trapped rodents are brought to the processing table contained within a trap following capture. The rodents are euthanised following recommended procedures with halothene and cervical dislocation by trained personnel. Kevlar gloves and personal protective equipment are used during this process. Rodents are placed on a processing mat where photos are obtained during the data collection process with the rodents unique identifier. Morphometric measurements are then recorded alongside biological samples for subsequent molecular identification to species level and assay for Lassa fever virus antibodies or acute infection. Rodent carcasses are then safely disposed of on site to reduce the risk of any infectious material remaining in the study environment. All data is collected in the field electronically using Open Data Kit questionnaires with photographic records collected as required. Samples are then returned to the study laboratory at Mercy Hospital, Bo, Sierra Leone where they remain in storage until analysis can be performed.

#### Locating trap sites

The GPS devices can be used to precisely locate the trapping grids within the villages. First turn on the device and press the find button on the front above mark. Use the direction buttons to select Coordinates and press enter. This takes to a screen that asks you to enter the location. The left and right arrows at the bottom of the screen move across the numbers. Select the value to change using the directional buttons and enter to select. This will then produce a purple line that will guide you to the coordinates. Tie some ribbon to a plant to identify this corner of the grid and then perform the same for the 3 other corners.

Coordinates below are given in degrees decimal minutes (DdM), this is consistent with the GPS devices used by the field team.

##### Baiama

We have set **six** trapping grids in Baiama.

1. **Grid 1**: Forested site
    - Point 1 = 7° 49.4867 N and 11° 15.0235 W
    - Point 2 = 7° 49.4867 N and 11° 15.0235 W
    - Point 3 = 7° 49.4708 N and 11° 15.0098 W
    - Point 4 = 7° 49.4840 N and 11° 15.0161 W
2. **Grid 2**: Fallow land, new rice field
    - Point 1 = 7° 49.8157 N and 11° 15.235 W
    - Point 2 = 7° 49.8086 N and 11° 15.2235 W
    - Point 3 = 7° 49.7903 N and 11° 15.2436 W
    - Point 4 = 7° 49.7885 N and 11° 15.2318 W
3. **Grid 3**: This site lies near the old Baiama village
    - Point 1 = 7° 49.8151 N and 11° 15.7421 W
    - Point 2 = 7° 49.8179 N and 11° 15.729 W
    - Point 3 = 7° 49.8245 N and 11° 15.7529 W
    - Point 4 = 7° 49.8386 N and 11° 15.7509 W
4. **Grid 4**: Banana plantation
    - Point 1 = 7° 50.1875 N and 11° 15.9803 W
    - Point 2 = 7° 50.1968 N and 11° 15.9904 W
    - Point 3 = 7° 50.1785 N and 11° 15.9953 W
    - Point 4 = 7° 50.1844 N and 11° 16.0042 W
5. **Grid 5**: Village, outside households
    - Point 1 = 7° 50.2663 N and 11° 16.132 W
    - Point 2 = 7° 50.2555 N and 11° 16.0783 W
    - Point 3 = 7° 50.18 N and 11° 16.043 W
    - Point 4 = 7° 50.171 N and 11° 16.093 W
6. **Grid 6**: Village, within households

- 4 traps per home

**Lalehun**

We have set **seven** trapping grids in Lalehun.

1. **Grid 1**: Edge of the village
    - Point 1 = 8° 11.801 N and 11° 4.767 W
    - Point 2 = 8° 11.782 N and 11° 4.742 W
    - Point 3 = 8° 11.781 N and 11° 4.775 W
    - Point 4 = 8° 11.769 N and 11° 4.758 W
2. **Grid 2**: Within and near a wet rice field
    - Point 1 = 8° 11.921 N and 11 4.771 W
    - Point 2 = 8° 11.94 N and 11 4.758 W
    - Point 3 = 8° 11.923 N and 11 4.727 W
    - Point 4 = 8° 11.908 and 11 4.739 W
3. **Grid 3**:
    - Point 1 = 8° 11.9417 N and 11 4.811 W
    - Point 2 = 8° 11.92 N and 11 4.822 W
    - Point 3 = 8° 11.967 N and 11 4.826 W
    - Point 4 = 8° 11.953 N and 11 4.838 W
4. **Grid 4**: Forest area
    - Point 1 = 8° 11.644 N and 11° 4.699 W
    - Point 2 = 8° 11.687 N and 11° 4.696 W
    - Point 3 = 8° 11.644 N and 11° 4.681 W
    - Point 4 = 8° 11.687 N and 11° 4.68 W
5. **Grid 5**: Cassava plantation
    - Point 1 = 8° 11.619 N and 11 4.811 W
    - Point 2 = 8° 11.633 N and 11 4.831 W
    - Point 3 = 8° 11.647 N and W 11 4.832 W
    - Point 4 = 8° 11.635 N and 11 4.806 W
6. **Grid 6**: Village, outside households
    - Point 1 = 8° 11.911 N and 11° 4.797 W
    - Point 2 = 8° 11.819 N and 11° 4.79 W
    - Point 3 = 8° 11.872 N and 11° 4.828 W
    - Point 4 = 8° 11.809 N and 11° 4.818 W
7. **Grid 7**: Village, within households
    - 4 traps per home

**Lambayama**

We have set **six** trapping grids in Lambayama.

1. **Grid 1**: Rice fields
    - Point 1 = 7° 51.042 N and 11° 11.739 W
    - Point 2 = 7° 51.038 N and 11° 11.7072 W
    - Point 3 = 7° 51.001 N and 11° 11.755 W
    - Point 4 = 7° 50.99 N and 11° 11.73 W

2. **Grid 2**: Cassava field
    - Point 1 = 7° 51.085 N and 11° 11.658 W
    - Point 2 = 7° 51.08 N and 11° 11.691 W
    - Point 3 = 7° 51.0579 N and 11° 11.6698 W
    - Point 4 = 7° 51.055 N and 11° 11.689 W
3. **Grid 3**: Fallow land
    - Point 1 = 7° 50.9661 N and 11° 11.555 W
    - Point 2 = 7° 50.963 N and 11° 11.53 W
    - Point 3 = 7° 50.947 N and 11° 11.535 W
    - Point 4 = 7° 50.949 N and 11° 11.557 W
4. **Grid 4**: Vegetable gardens
    - Point 1 = 7° 50.9958 N and 11° 11.8425 W
    - Point 2 = 7° 50.9989 N and 11° 11.8126 W
    - Point 3 = 7° 50.979 N and 11° 11.792 W
    - Point 4 = 7° 50.9659 N and 11° 11.8175 W
5. **Grid 5**: Village, outside households
    - Point 1 = 7° 51.064 N and 11° 11.741 W
    - Point 2 = 7° 50.9901 N and 11° 11.7869 W
    - Point 3 = 7° 51.018 N and 11° 11.824 W
    - Point 4 = 7° 51.0953 N and 11° 11.7993 W
6. **Grid 6**: Village, within households
    - 4 traps per home

**Seilama**

We have set up **six** sites in Seilama.

1. **Grid 1**: Palm plantation, near the village and main road
    - Point 1 = 8° 7.33498 N and 11° 11.5399 W
    - Point 2 = 8° 7.318 N and 11° 11.531 W
    - Point 3 = 8° 7.3181 N and 11° 11.5578 W
    - Point 4 = 8° 7.338 N and 11° 11.568 W
2. **Grid 2**: Cacao and Coffee plantation
    - Point 1 = 8° 7.378 N and 11° 11.649 W
    - Point 2 = 8° 7.4 N and 11° 11.643 W
    - Point 3 = 8° 7.413 N and 11° 11.653 W
    - Point 4 = 8° 7.384 N and 11° 11.67 W
3. **Grid 3**: Dry rice field
    - Point 1 = 8° 7.424 N and 11° 11.657 W
    - Point 2 = 8° 7.446 N and 11° 11.66 W
    - Point 3 = 8° 7.467 N and 11° 11.672 W
    - Point 4 = 8° 7.443 N and 11° 11.685 W
4. **Grid 4**: Wet rice plantation
    - Point 1 = 8° 7.2507 N and 11° 11.6773 W
    - Point 2 = 8° 7.2568 N and 11° 11.6191 W
    - Point 3 = 8° 7.2287 N and 11° 11.6747 W
    - Point 4 = 8° 7.2376 N and 11° 11.6266 W

5. **Grid 5**: Forest
    - Point 1 = 8° 7.441 N and 11° 11.84 W
    - Point 2 = 8° 7.4434 N and 11° 11.8737 W
    - Point 3 = 8° 7.422 N and 11° 11.89 W
    - Point 4 = 8° 7.4149 N and 11° 11.869 W
6. **Grid 6**: Village, outside households
    - Point 1 = 8° 7.3151 N and 11° 11.5978 W
    - Point 2 = 8° 7.315 N and 11° 11.64 W
    - Point 3 = 8° 7.3756 N and 11° 11.6048 W
    - Point 4 = 8° 7.3741 N and 11° 11.6334 W
7. **Grid 7**: Village, within households
    - 4 traps per home

Grids 6 and 7 will be combined for subsequent statistical analysis as rodents are expected to move between households and in the areas surrounding houses.

#### Data collection

Three questionnaires have been developed in the ODK app for data collection. The first, trap setup must be completed during the process of setting the traps as we collect information on the location of each trap. The second, trap check, is completed on each morning of the trapping session to record whether a rodent was trapped. The third, rodent sampling, is conducted alongside the measurement and biopsy of each individual.

It is advised that a single field worker is responsible for data input alongside the person or people placing traps, checking traps and conducting biopsies. If there are any difficulties with data entry onto the digital devices paper versions of the questionnaires are available.

##### Trap setup

Mark the extent of the trapping grid by placing ribbons at the locations of the above coordinates. Place traps in a 7 by 7 grid spaced 10 metres apart within the grid. For each trap record its coordinates in the ODK questionnaire on the mobile phone app. If required take a photo of the trap location if you are unable to decide the specific habitat type of the trap. Place bait in the trap and close it. Once all traps have been set and recorded the traps can be opened at dusk for the first trap night. During the trap check in the morning, traps are closed for the day prior to refreshing the bait. On each subsequent day the bait is refreshed with traps opened again at dusk.

##### Trap check

Each morning every trap is checked for the presence/absence of bait, whether the trap has snapped shut over night and whether a rodent has been trapped. The weather overnight is also collected on this data entry form on the ODK app.

#### Rodent sampling

Trapped rodents are located at the trap check. The traps are placed in plastic bags and brought to the autopsy site. Rodents are euthanised prior to morphological measurements and sampling. Processes for rodent handling and measurements were conducted as outlined in previously published guidance (Mills et al. 1995; Fichet-Calvet 2014).

1. Identification
    - Rodent unique number
    - Trap number

- Trap night
- Initial species/genus identification

2. Morphological measures
    - Weight in grams
    - Length of head to base of tail (head body) in mm
    - Length of the tail in (note whether tail is cut) mm
    - Length of the hind foot (not including claws) in mm
    - Diameter of the ear measured from the pinna to the edge of the ear in mm
    - Length of the skull from the occiput to the tip of the nose, for shrews measure to the end of the projecting teeth in mm
3. Autopsy measures
    - Rodent sex (M/F)
    - Presence of internal or external testes for males
    - Development of seminal vesicles for males
    - The identification of a perforate vagina for females
    - The presence of visible teats for females and the number of pairs of nipples
    - The number of developing embryos for females
4. Sample collection
    - Document whether the following samples have been successfully obtained
        - Photo of rodent
        - Serum sample, in vial and on filter paper
        - Tissue sample of liver and spleen
        - Tissue sample of ear
        - Eye of rodent

The first step of rodent sampling is to set up the biopsy table, clean the area and ensure adequate personal protective equipment is available. Assign one team member to perform the rodent euthanisation. Assign another to conduct real-time data entry. Two team members are required for the biopsy process.

First, rodents are euthanised and assigned a unique identifier based on the study visit, the study village and the order in which they are being processed. This is written on a piece of paper and photographed alongside a dorsal (place the rodent belly down) and ventral (place the rodent belly up) image of the rodent. The trap number the rodent was found in is and the trap night is then recorded. The field identification of the species is then recorded, the identification key is used for reference (Rodent Identification), this is based on two published taxonomic keys (Happold and Happold 2013; Monadjem et al. 2015). Next we obtain external measurements of the rodents.

Samples are placed in labelled cryovials. Formalin is added to eye and skin tag samples to aid preservation. Ethanol is added to liver and spleen samples.

#### Data collection details

##### Direct ODK entry

There are three forms you can access through ODK connect on your mobile phone or the study team tablets. The forms, once saved, will automatically be sent to the ODK server once they can connect to the internet. There is a sim card in the tablet that can be loaded with credit.

1. site_setup_v2: This sheet is completed for each site on the first day of trapping. It is important to ensure you correctly write the trap number and it's coordinates. If you make any errors you can edit the file or notify Dianah/David and they can amend it. You will describe each site, the habitat and surroundings of each trap and the coordinates for each trap. Photos can be taken if you are having difficulty completing the questions.

2. trap_check_v1: This sheet is used to collect information about the number of traps missing bait, if they have been sprung shut or if they contain rodents the next morning. It may be easier to note the traps on a piece of paper first and then to enter the data into ODK.
3. rodent_v1: This sheet is used to collect information about the trapped rodent. Ensure that the trap number and rodent number are correct. The trap number is important to know where the rodent came from. The rodent number should be made by putting the number of the visit, then the 3 letters of the village and then the number this rodent is for this visit.

For example:

- The 12th rodent trapped in the 2nd visit in Seilama would be 2SEI-012
- The 3rd rodent trapped on the 1st visit in Baiama would be 1BAI-003

##### Sample inventory

Following field collection all samples will be checked into the laboratory on the sample inventory sheet. Here the labelling on the samples will be checked and the samples will be stored in the same container for village study site and visit. The data from the sample inventory will be used to cross check against the ODK entered samples to ensure consistency of labelling and traceability.

##### Sample storage

Samples are collected in sterile cryovials. Eyes and ear snips are stored in Formalin, while liver and spleen samples are stored in Ethanol. Whole blood is aspirated onto filter papers which are stored in single use bags with dessicant. 1-2ml of whole blood is stored in a cryovial.The rodent identifier is written onto the cryovials in indelible ink and they are collected by sample type for each study village and visit in containers. At the end of each day samples are transferred into field freezers for storage prior to transport to -20°C storage at the laboratory.

### Laboratory protocols

#### DNA extraction

This is the protocol to identify rodents to species level using PCR for use in Sierra Leone.

Three kits are required in addition to consumables for each run.

1. QIAgen DNeasy Blood and Tissue kit. This protocol will be performed on spleen or liver tissue. However, in case of issues purchase an additional buffer AL as this protocol can also be run on blood samples as long as AL hasn't been pre-diluted with ethanol. This step extracts DNA from the sample.
2. Invitrogen Platinum *Taq* DNA Polymerase. This is used to produce the mastermix that will amplify the DNA bound to the primers.
3. Primers, obtained from Eurofins (Bangura et al. 2021):
   - *M. natalensis* specific primer
     1. F-607: 5' CGG GCT CTA ATA ACC CAA CG 3'
     2. R-813: 5' TTC TGG TTT GAT ATG GGG AGG T 3'
   - *M. erythroleucus* specific primer
     1. F-49: 5' CAT TCA TTG ACC TAC CTG CT 3'
     2. R-505: 5' AGA ATC CCC CTC AAA TTC AC 3'
   - Rodent *CytB* for subsequent sequencing
     1. L-7: 5' ACC AAT GAC ATG AAA AAT CAT CGT T 3'
     2. H-15915: 5' TCT CCA TTT CTG GTT TAC AAG AC 3'

###### Kit contents

**QIAgen DNeasy 250 samples**

| Component | Supplied amount/packaging | Storage requirements |
|---|---|---|
| DNeasy mini spin columns | 250 tubes | room temperature |
| Collection tubes | 500 tubes | room temperature |
| Buffer ATL | 50ml | room temperature |
| Buffer AL | 2 x 33ml | room temperature |
| Buffer AW1 (concentrate) | 98ml | room temperature |
| Buffer AW2 (concentrate) | 66ml | room temperature |
| Buffer AE | 2 x 60ml | room temperature |
| Proteinase K | 6ml | room temperature |

Additional consumables Pipette tips, microcentrifuge tubes 1.5ml, chlorine tablets, PBS tablets, 96-100% EtOH.

Additional equipment Pipettes, scalpel, tile, tweezers, microcentrifuge, thermomixer, vortex.

###### Invitrogen Platinum *Taq* DNA Polymerase 600 reactions

| Component | Supplied amount/packaging | Storage requirements |
|---|---|---|
| Platinum *Taq* DNA polymerase | 120µl | -20°C (freezer) |
| 10x PCR buffer | 3 x 1.2ml | -20°C (freezer) |
| 50mM Magnesium Chloride | 1ml | -20°C (freezer) |

Additional reagents

| Component | Supplied amount/packaging | Storage requirements |
|---|---|---|
| dNTP mix | 100µl | -20°C (freezer) |
| nuclease free water | 10 x 1.5ml | -20°C (freezer) |
| DNA ladder | 250µg | -20°C (freezer) |
| Midori green agarose | 100 tablets | room temperature |
| TAE buffer | 50x concentrate | -4°C (fridge) |
| DNA loading dye | 5 x 1ml | room temperature |

Additional consumables Pipette tips, microcentrifuge tubes 1.5ml, PCR tubes.

Additional equipment Pipettes, microcentrifuge, thermomixer, vortex, PCR cycler, gel electrophloresis tray, gel combs, gel reader.

###### Primers

| Component | Supplied amount/packaging | Storage requirements |
|---|---|---|
| *M. natalensis* specific primer | NA | -20°C (freezer) |
| *M. erythroleucus* specific primer | NA | -20°C (freezer) |
| Rodent *CytB* | NA | -20°C (freezer) |

Additional reagents Nuclease free water for dilution to working solution

###### Protocol

The majority of this protocol will be conducted in Sierra Leone, unfortunately due to no available sequencing in country purified, stable, DNA products will be transferred to Germany for commercial sequencing.

**Sample selection** Samples will be batch processed. Each gel can contain 14 rodent samples, a negative extract and a DNA ladder. All samples should undergo DNA extraction, the subsequent testing will need to be selected based on the field identification. We will be performing confirmatory testing on all identified *Mastomys sp.* and testing for all individuals that may be confused with this species.

Species identified in the field as *Crocidura sp.*, *Lophuromys sp.* and *Lemniscomys sp.* do not need testing with *Mastomys* specific primers. For other samples they will first be tested with *M. natalensis* specific primers, if these are positive those samples do not need to be tested further. All remaining samples will be tested with *M. erythroleucus* specific primers. Those that are positive for either of the *Mastomys* primers will be stored but do not need further testing. All samples except those identified as *M. natalensis* or *M. erythroleucus* will then be processed for rodent specific *CytB* for subsequent sequencing. Once *CytB* PCR amplification has been confirmed through gel electrophoresis these samples can be stored for transfer to the UK. Any samples that failed to produce a single band confirmatory for *CytB* amplification will need to be re-processed.

The same DNA extract if there is no evidence of cross contamination can be used for all three PCR reactions.

##### DNA extraction

1. Remove samples from the freezer, organise them and ensure that each is uniquely labelled.
2. If formalin fixed tissue is being used rinse the sample twice in PBS to remove the fixative.
3. Cut up to 10mg of tissue into small pieces on a clean tile with clean instruments, place into a labelled microcentrifuge tube.
4. Decontaminate equipment and tile, repeat for all samples being processed.
5. Add 180µl of buffer ATL to each tube, ensure that samples are covered by liquid reagent.
6. Add 20µl of proteinase K to each vial and vortex thoroughly.
7. Place tubes into a thermomixer at 56°C until the tissue is completely lysed ~ 1-3 hours.
8. Once lysed vortext the samples
9. Prepare a 1:1 mix of buffer AL and ethanol 96-100% and vortex. 400µl of reagent are added to each sample tube so calculate the volumes required prior to mixing AL and ethanol. For example 14 sample and a negative extract will require 6ml of the combined solution so make up 6.2ml
10. Vortex tubes after addition of the combined AL/ethanol.
11. A gelatinous layer may form, ensure adequate mixing by shaking the vials vigorously.
12. Pipette this mixture into a labelled spin column. Set the pipette to withdraw 620µl to ensure all volume is withdrawn.
13. Centrifuge at 6,000g (8000rpm) for 1 minute in the spin column and discarge the collection tube.
14. Place the spin column in a new, clean collection tube.
15. Add 500µl of buffer AW1 and centrifuge at 6,000g (8000rpm) for 1 minute. The flow through can be discarded and the collection tube re-used.
16. Label 1.5ml microcentrifuge tubes according to the samples in preparation for step 17.
17. Add 500µl of buffer AW2 and centrifuge at 20,000g (14000rpm) for 3 minute. The flow through can be discarded, ensure that there is no contact between the flow through and the membrane as the purpose of this step is to dry the membrane. If contact occurse, empty the collection tube and centrifuge again at 20,000g (14000rpm) for 1 minute. Place spin column in corresponding microcentrifuge tube.
18. Pipette 200µl buffer AE directly onto the membrane and incubate for 1 minute.
19. Centrifuge at 6,000g (8000rpm) for 1 minute. The spin column can now be discarded, the labelled microcentrifuge tube now contains the extracted DNA.
20. Place tubes into a sealed bag with a label for the date of extraction. These tubes can be stored in the freezer for subsequent PCR and downstream sequencing.

##### Preparing mastermix and combine with samples

1. An excel document for each of the primers is available to calculate the proportions of reagents in the mastermix. Print this out after entering the number of samples being run for each primer set.
2. Take out the DNA extract samples from the freezer.
3. Take out the *Taq* DNA Polymerase kit from the freezer.
4. Take out the primers from the freezer.

5. Produce a working dilution of the relevant primers according to manufacturers guidance in 2 microcentrifuge tubes, 1 for each primer direction. Dilute using DNAse free water.
6. Combine the reagents for the mastermix as guided by the excel sheet. Ensure tube is vortexed and then centrifuged for good mixing.
7. Place the PCR tubes into the cold PCR tube holder.
8. Place the required volume of the mastermix into each PCR tube, the same pipette tip can be used for this.
9. Arrange samples as they will be placed in the PCR tubes.
10. Place the required volume of sample into each PCR tube. The pipette tip needs to be changed between samples.
11. Close the tube lids as you add in sample to ensure you don't wrongly combine samples.
12. Label the tubes as per the earlier arrangement.

##### PCR amplification

1. Set up the cycler as per the temperature profile guidance in the excel spreadsheet or below.
2. Spin PCR tubes to ensure good mixture of sample and mastermix.
3. Place tubes in the cycler and run the programme.
4. This will take around 2-3 hours.

| PCR method | Temperature | Time | Cycles |
|---|---|---|---|
| *M natalensis* | 94°C | 3 minutes | - |
| - | 94°C | 30 seconds | x35 |
| - | 55°C | 30 seconds | x35 |
| - | 72°C | 30 seconds | x35 |
| - | 72°C | 10 minutes | - |
| - | -4°C | ongoing | - |
| *M erythroleucus* | 94°C | 3 minutes | - |
| - | 94°C | 30 seconds | x35 |
| - | 55°C | 30 seconds | x35 |
| - | 72°C | 45 seconds | x35 |
| - | 72°C | 10 minutes | - |
| - | -4°C | ongoing | - |
| *CytB* | 95°C | 3 minutes | - |
| - | 94°C | 30 seconds | x35 |
| - | 52°C | 40 seconds | x35 |
| - | 72°C | 90 seconds | x35 |
| - | -4°C | ongoing | - |

##### Gel preparation

1. We will use Midori green agarose to identify locations of DNA bands.
2. Each gel is made up of 2% agarose and 30ml is required to provide a 2 row gel.
3. Dissolve get tablets in water and heat if required to ensure crystals dissolved.
4. Pour 30ml into a gel tray.
5. Allow to set ~ 30mins, remove combs and can then be stored in TAE in the fridge until required.

##### Run gel

1. Arrange samples in PCR tubes in the order they will be loaded on the gel.
2. Place 4µl of the DNA ladder in the first well.
3. Place ~1µl of loading dye onto a paraffin sheet.
4. Pipette 4µl of sample into the loading dye bubble, withdraw to mix the dye and sample and place directly into the well. Change pipette tips between samples.

5. In the last well place the negative extract control.
6. Once gel is loaded place into an electrophoresis bath and set to 80 V, 400 mA and run for 40 minutes.

**Interpreting gel**

1. Take gel to the reader.
2. Take a photo of the gel, interpret as follows.
    1. *M. natalensis* specific primer: Bands seen should all align along a similar mass on the ladder. If this is the case the samples have been adequately amplified with good specificity. In addition the negative extract must be negative. Together these results will mean the gel is interpretable. If the negative extract shows a band this is indicative of contamination at an earlier point of the process and the DNA extraction would need to be repeated for these samples. Document the results of the gel as follows:
        - Species confirmed as *M. natalensis*
        - Species confirmed as not *M. natalensis*
    2. *M. erythroleucus* specific primer: Bands seen should all align along a similar mass on the ladder. If this is the case the samples have been adequately amplified with good specificity. In addition the negative extract must be negative. Together these results will mean the gel is interpretable. If the negative extract shows a band this is indicative of contamination at an earlier point of the process and the DNA extraction would need to be repeated for these samples. Document the results of the gel as follows:
        - Species confirmed as *M. erythroleucus*
        - Species confirmed as not *M. erythrolecus*
    3. *CytB* specific primer: Bands seen should all align along a similar mass on the ladder. If this is the case the samples have been adequately amplified with good specificity. In addition the negative extract must be negative. Together these results will mean the gel is interpretable. If the negative extract shows a band this is indicative of contamination at an earlier point of the process and the DNA extraction would need to be repeated for these samples. Document the results of the gel as follows:
        - Adequate *CytB* amplification, suitable for sequencing
        - Inadequate *CytB* amplification, not-suitable for sequencing. Needs to be repeated potentially starting from DNA extraction stage.

**Preparing samples for sequencing**

1. Samples with confirmed *CytB* can now be transferred to Germany for sequencing.
2. Samples can be transported at room temperature, in safelock sample tubes as DNA products are stable and non-infectious.
3. For the first transfer we will trial adding the PCR primers into the products in Sierra Leone and compare to the results when PCR primers are added in the UK.

**Sequence processing**

Eurofins provide a FASTA file via email for all attempted sequences. These are downloaded and saved for analysis using NCBI BLAST.

1. Log in to BLAST
2. Select Nucleotide BLAST
3. The FASTA file can be copied into the query sequence box, BLAST will recognise the number of distinct samples
4. Enter a descriptive title for the job, i.e., the number of sequences and date of file
5. Set the following for the search set
    - Standard databases, nucleotide collection
    - Organism; Rodentia (taxid: 9989) AND Soricomorpha (taxid: 9362)
6. Program selection is set as highly similar sequences (megablast)

7. Algorithm parameters
    - Max target sequences 100
    - Short queries selected
    - Expect threshold 0.05
    - Word size 28
    - Scoring parameters, match/mismatch scores 1,-2, gap costs linear
    - Filter low complexity regions selected and mask for lookup table only
8. Run BLAST
9. Complete the `sequencing.csv` record by entering the `rodent_uid` and PCR_id associated with the Eurofins barcode and enter the rodent species identified by BLAST
10. For failed sequences the program selection can be changed to somewhat similar sequences (blastn)
11. Failed sequences at this stage will need to be repeated

Mills, James N, James E Childs, Tom G Ksiazek, Clarence James Peters, and Wallis M Velleca. 1995. "Methods for Trapping and Sampling Small Mammals for Virologic Testing."

Monadjem, Ara, Peter J. Taylor, Christiane Denys, and Fenton P. D. Cotterill. 2015. *Rodents of Sub-Saharan Africa: A Biogeographic and Taxonomic Synthesis*. Berlin, München, Boston: DE GRUYTER. https://doi.org/10.1515/9783110301915.

### Supplementary Information

#### Supplementary Information 1

### Supplementary Figure 1

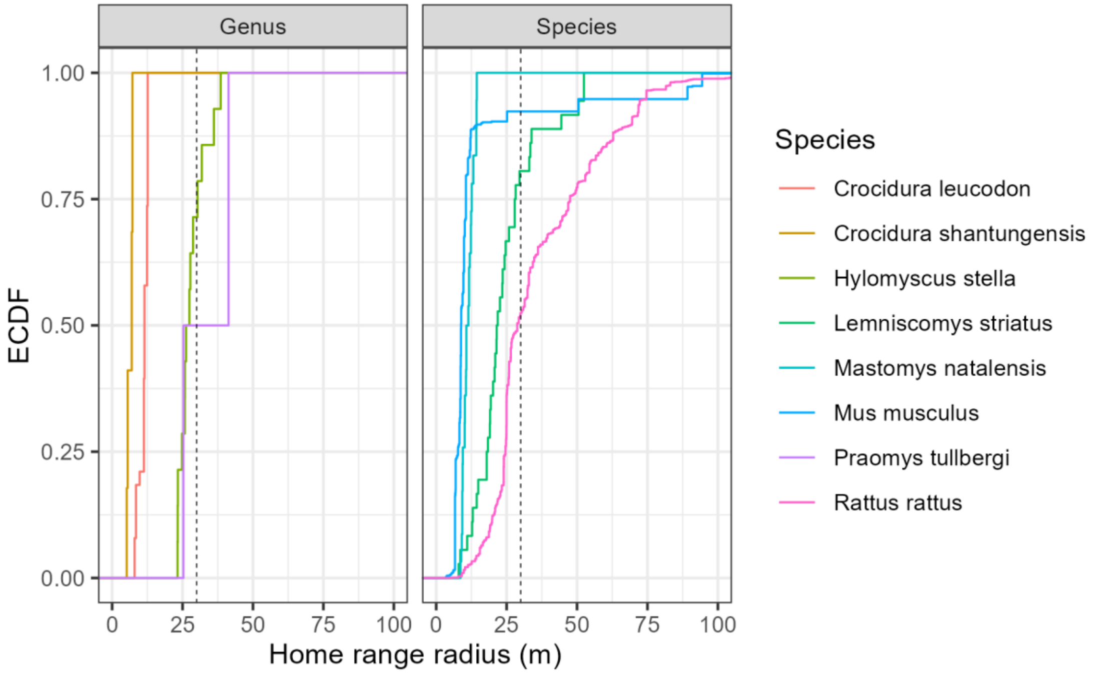

*Supplementary Figure 1: Empirical Cumulative Density Function of the home range radius of rodent and shrew species with data available in the HomeRange dataset. Species that match detected genera in our study include two shrew species Crocidura leucodon and Crocidura shantungensis and two rodent species Hylomyscus stella and Praomys tullbergi. Four species matches to rodent species detected in our study were also included Lemniscomys striatus, Mastomys natalensis, Mus musculus and Rattus rattus. Only Lemniscomys striatus and Mastomys natalensis contain data from Africa (Uganda and Tanzania respectively). The dashed line represents the 30m range radius used in one of the three sensitivity analyses in the current study.*

### Supplementary Figure 2

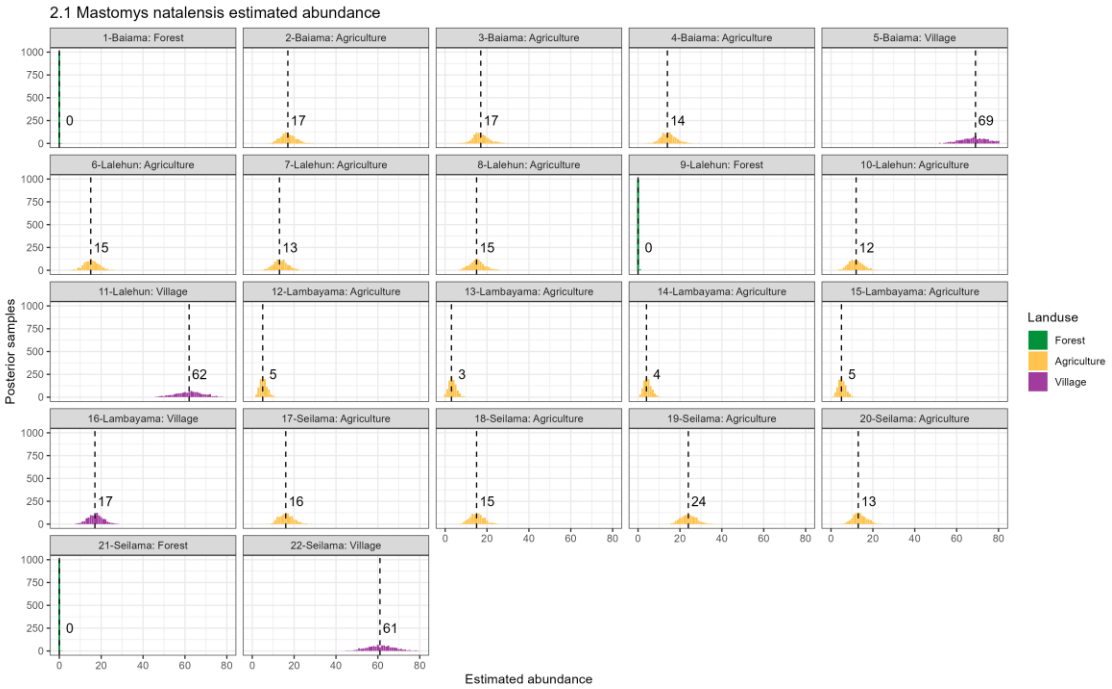

*Supplementary Figure 2.1: Estimated abundance at each sampling site for Mastomys natalensis. The dashed line and number is the median abundance used to infer the population size at this study site.*

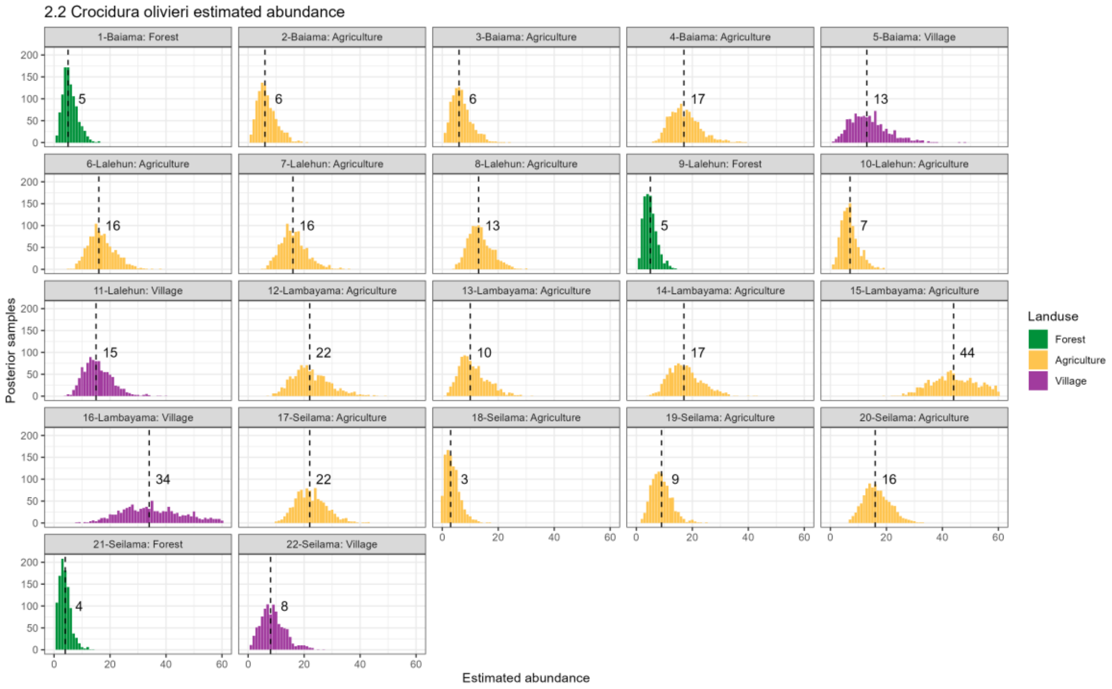

*Supplementary Figure 2.2: Estimated abundance at each sampling site for Crocidura olivieri. The dashed line and number is the median abundance used to infer the population size at this study site.*

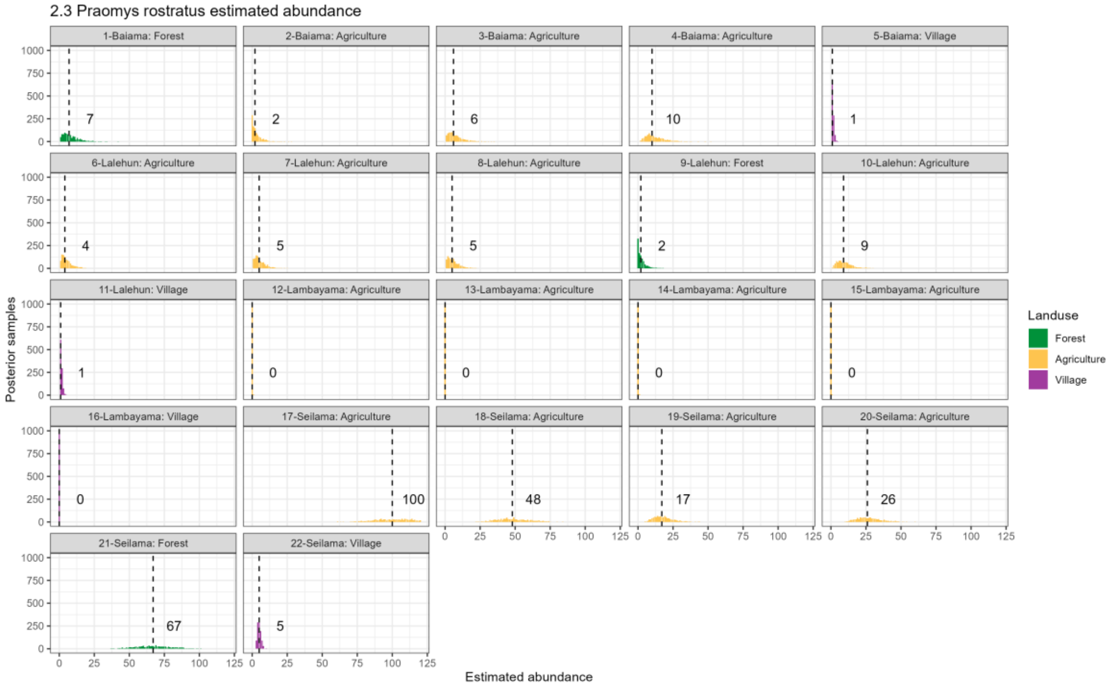

*Supplementary Figure 2,3: Estimated abundance at each sampling site for Praomys rostratus. The dashed line and number is the median abundance used to infer the population size at this study site.*

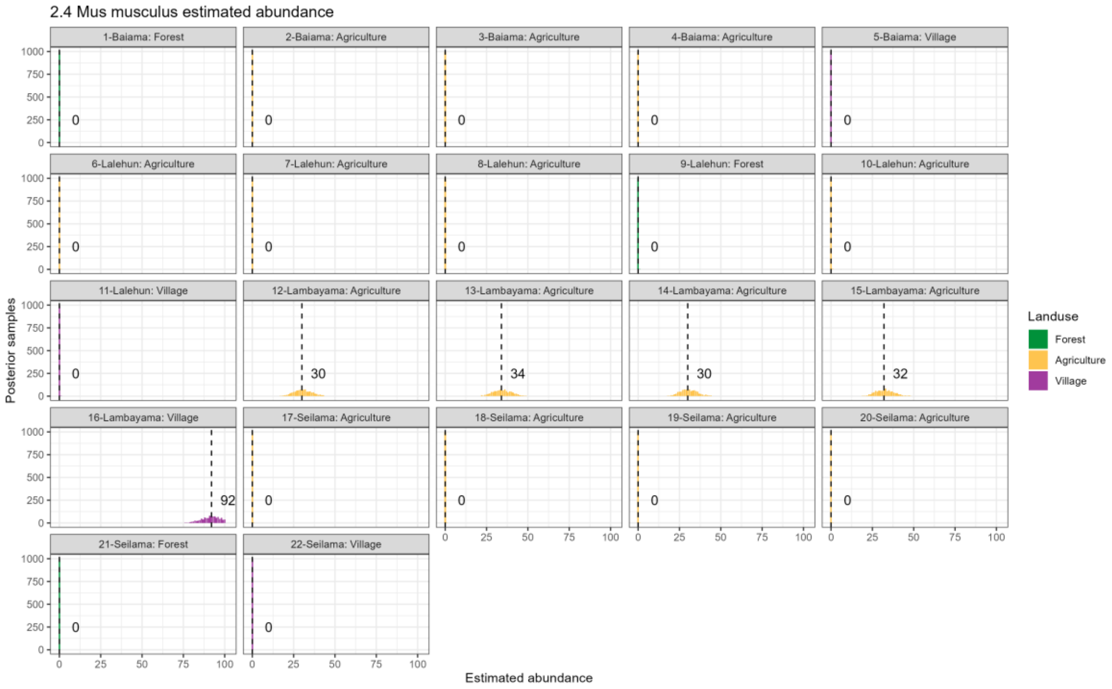

*Supplementary Figure 2.4: Estimated abundance at each sampling site for Mus musculus. The dashed line and number is the median abundance used to infer the population size at this study site.*

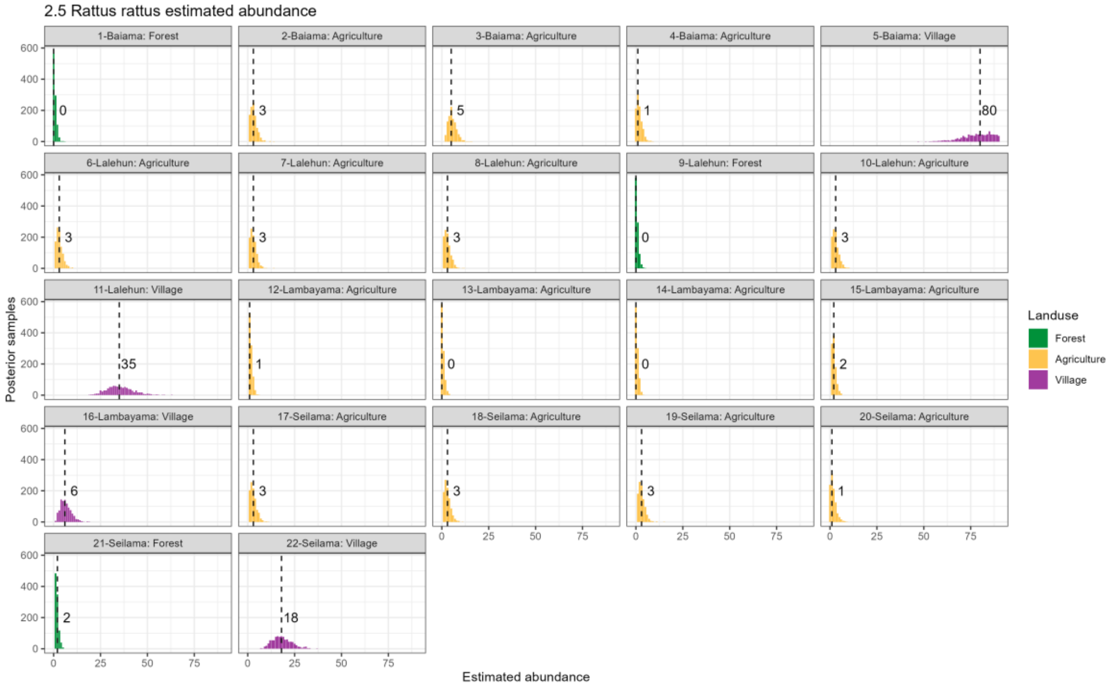

*Supplementary Figure 2.5: Estimated abundance at each sampling site for Rattus rattus. The dashed line and number is the median abundance used to infer the population size at this study site.*

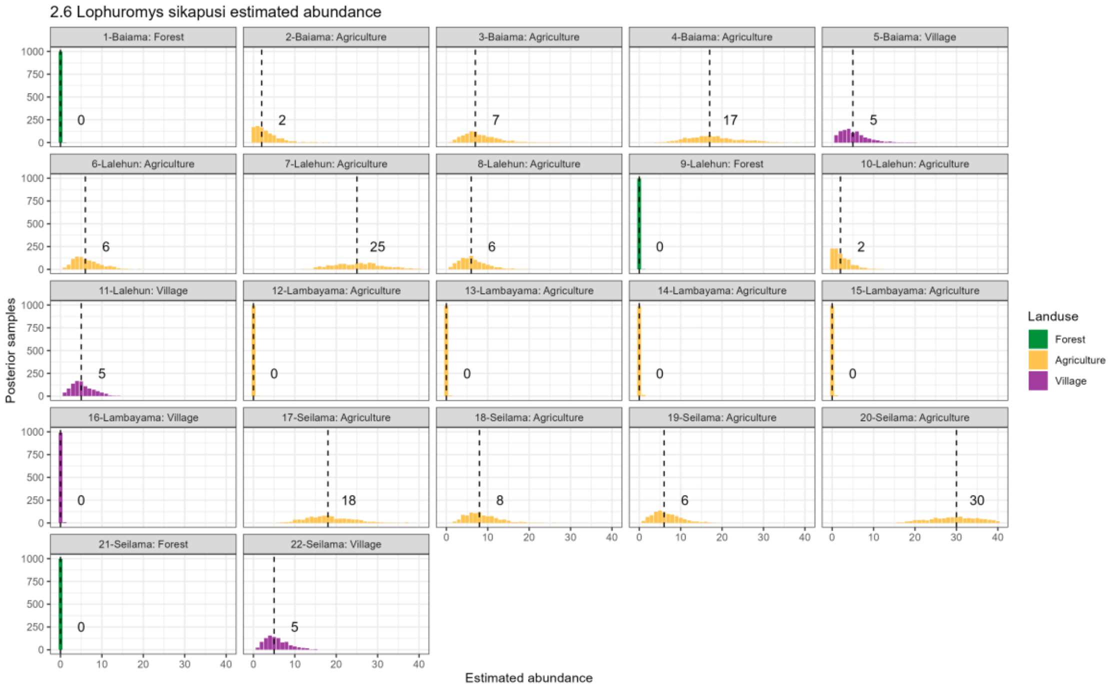

*Supplementary Figure 2.6: Estimated abundance at each sampling site for Lophuromys sikapusi. The dashed line and number is the median abundance used to infer the population size at this study site.*

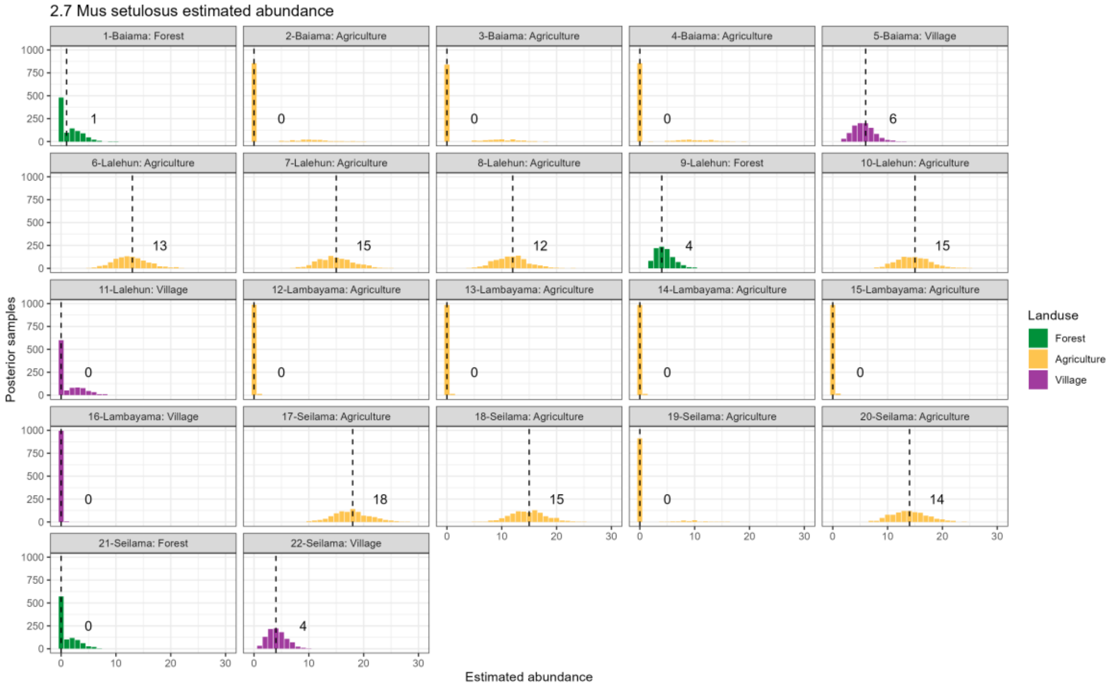

*Supplementary Figure 2.7: Estimated abundance at each sampling site for Mus setulosus. The dashed line and number is the median abundance used to infer the population size at this study site.*

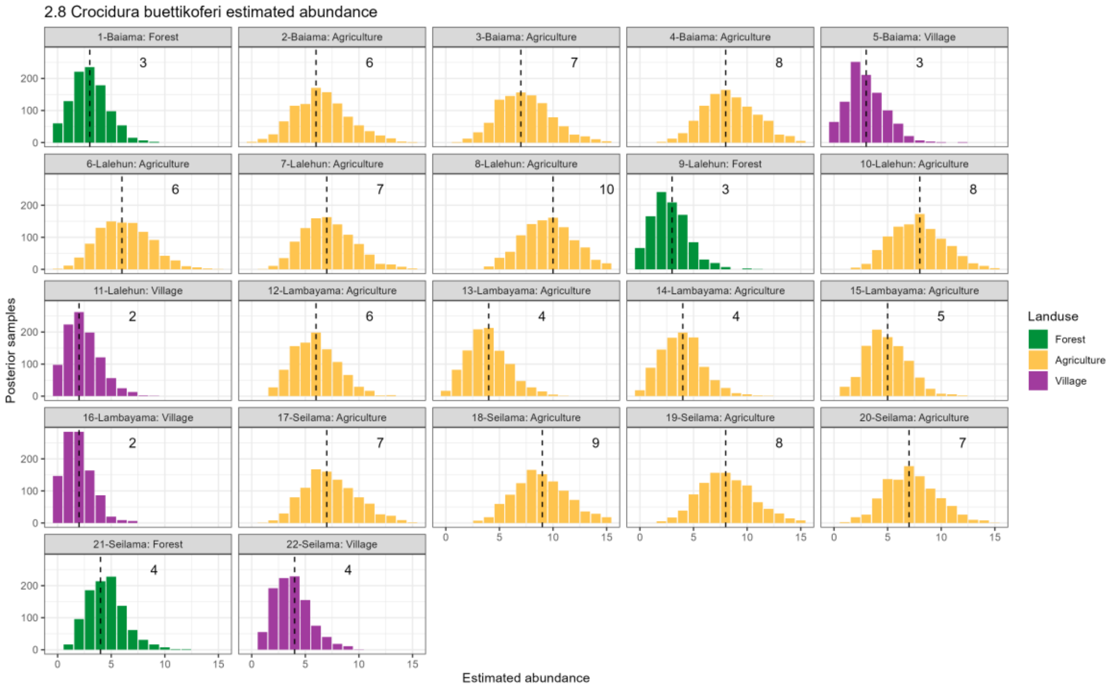

*Supplementary Figure 2.8: Estimated abundance at each sampling site for Crocidura buettikoferi. The dashed line and number is the median abundance used to infer the population size at this study site.*

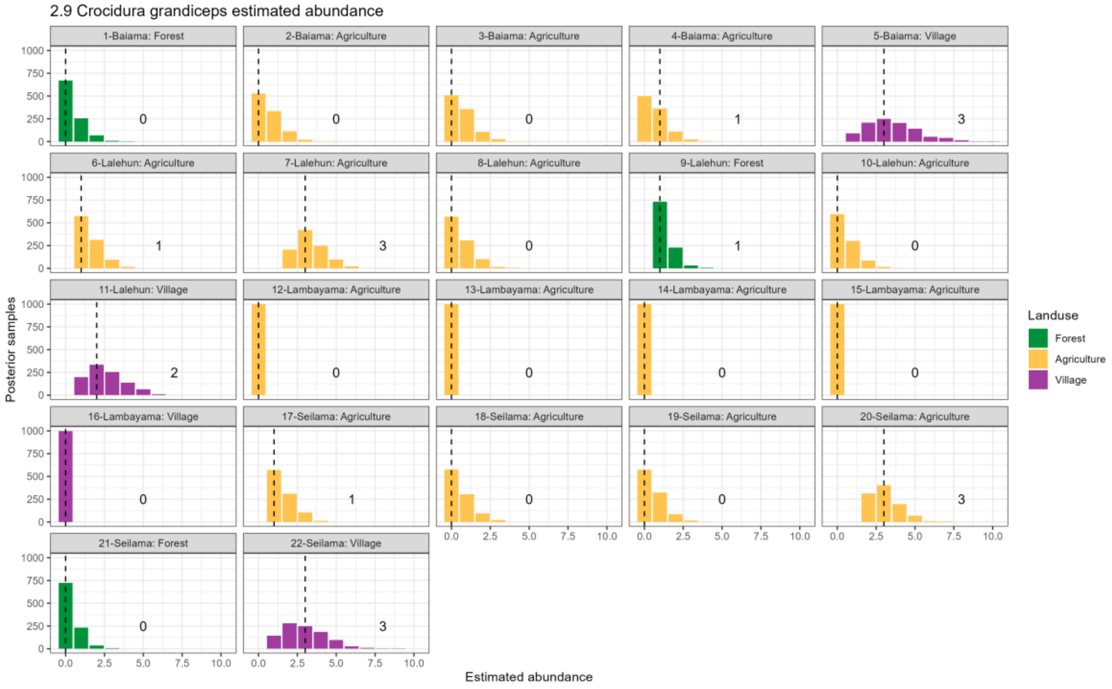

*Supplementary Figure 2.9: Estimated abundance at each sampling site for Crocidura grandiceps. The dashed line and number is the median abundance used to infer the population size at this study site.*

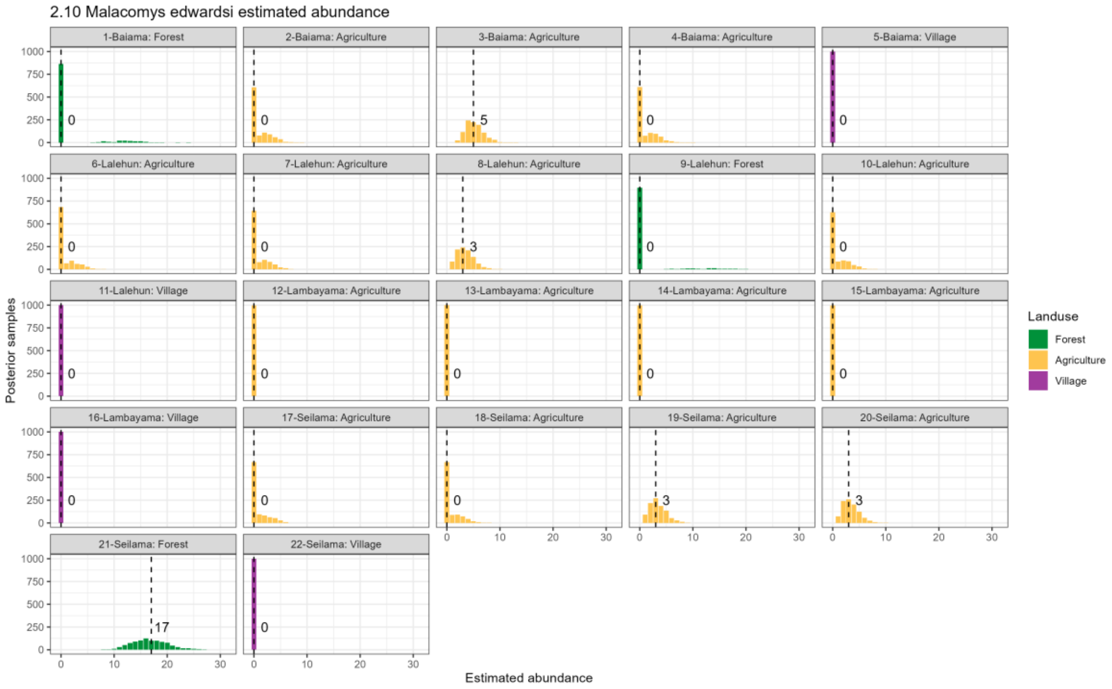

*Supplementary Figure 2.10: Estimated abundance at each sampling site for Malacomys edwardsi. The dashed line and number is the median abundance used to infer the population size at this study site.*

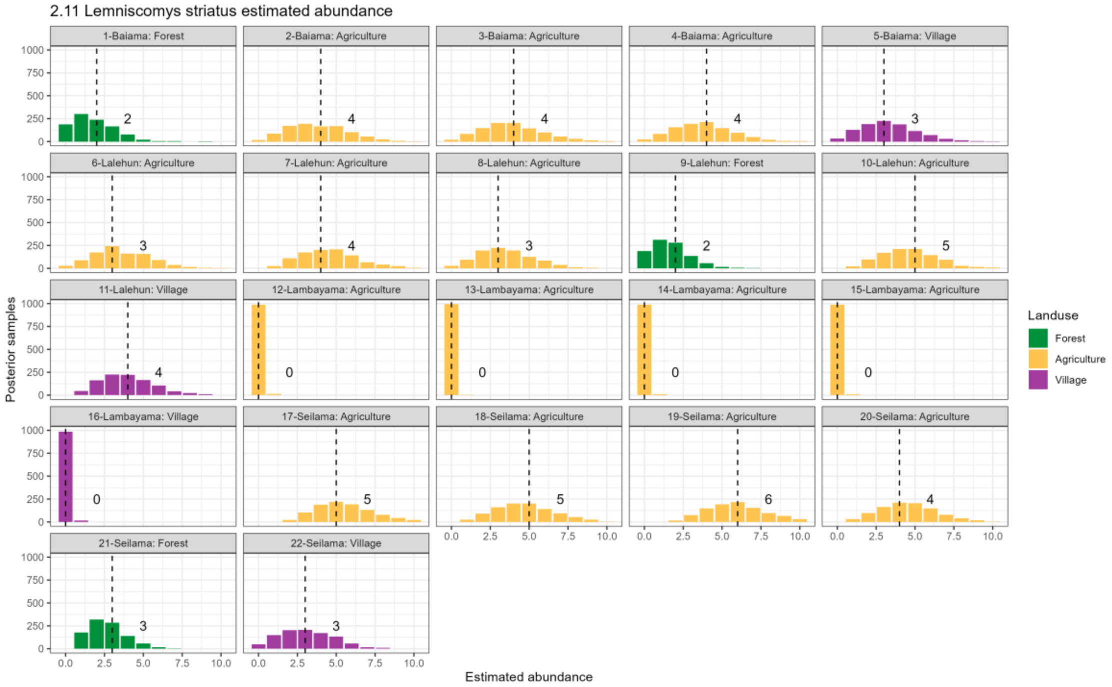

*Supplementary Figure 2.11: Estimated abundance at each sampling site for Lemniscomys striatus. The dashed line and number is the median abundance used to infer the population size at this study site.*

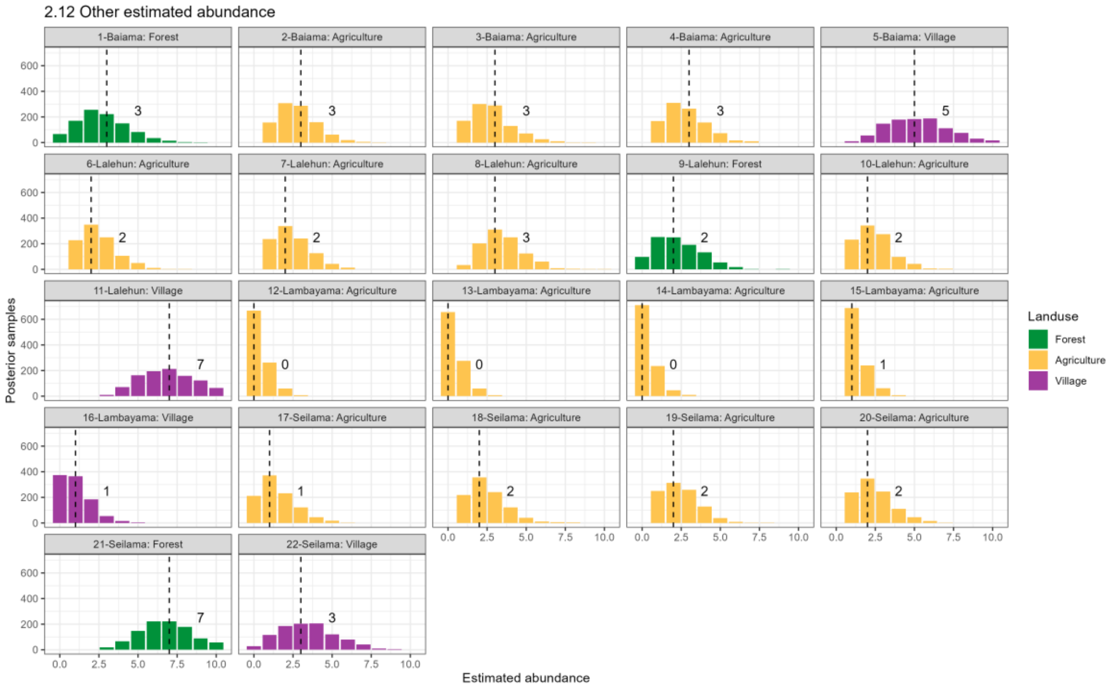

*Supplementary Figure 2.12: Estimated abundance at each sampling site for Other species. The dashed line and number is the median abundance used to infer the population size at this study site.*

### Supplementary Figure 3

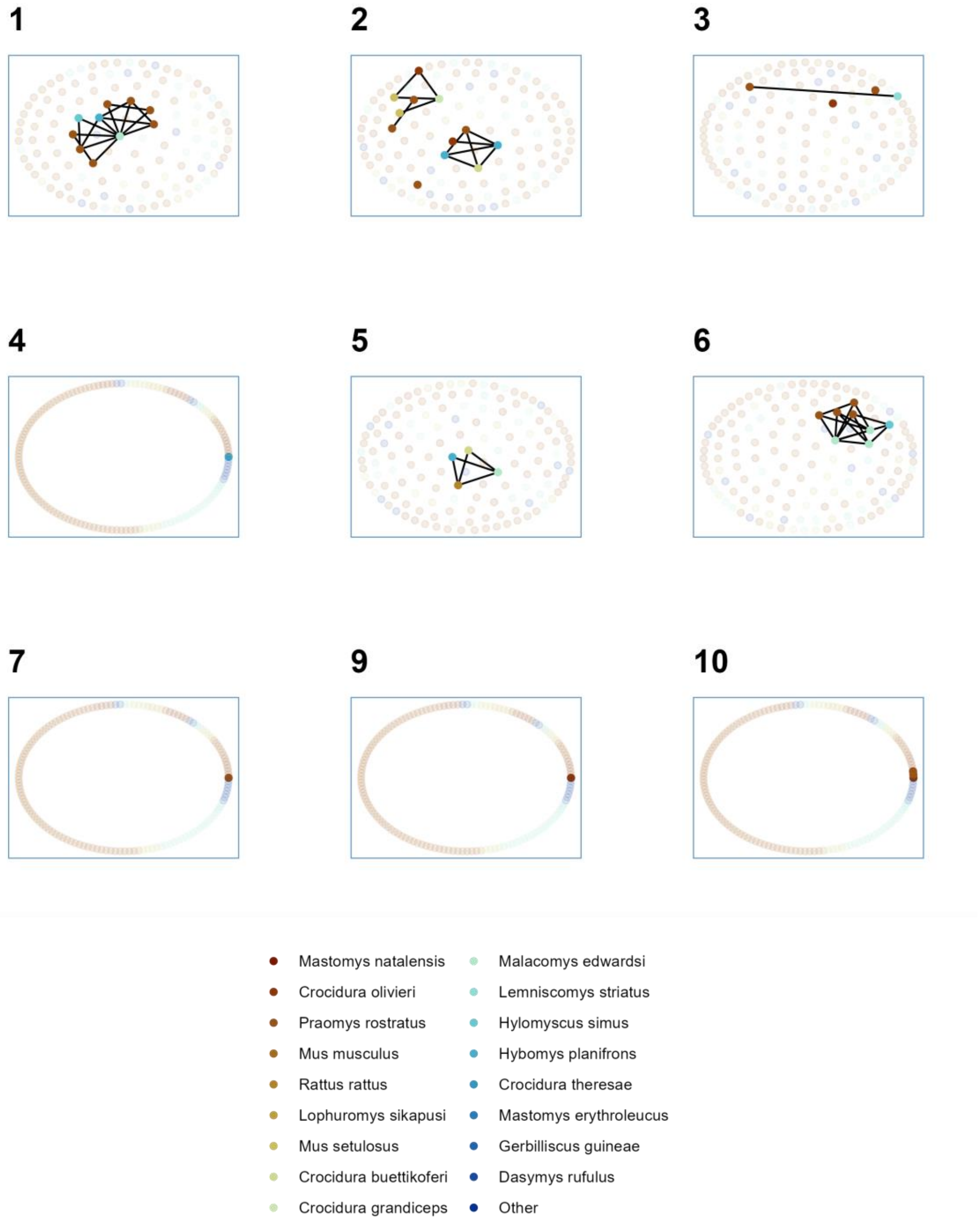

*Supplementary Figure 3.1: Networks produced from trapping sessions conducted in forest settings. Numbers refer to the trapping session, no individuals were detected in forest during trapping session 8. Observed individuals are shown in solid colours with colour referring to species. Unobserved individuals estimated from abundance modelling are shown in pale colours. Contacts between individuals (edges) are shown for observed individuals only.*

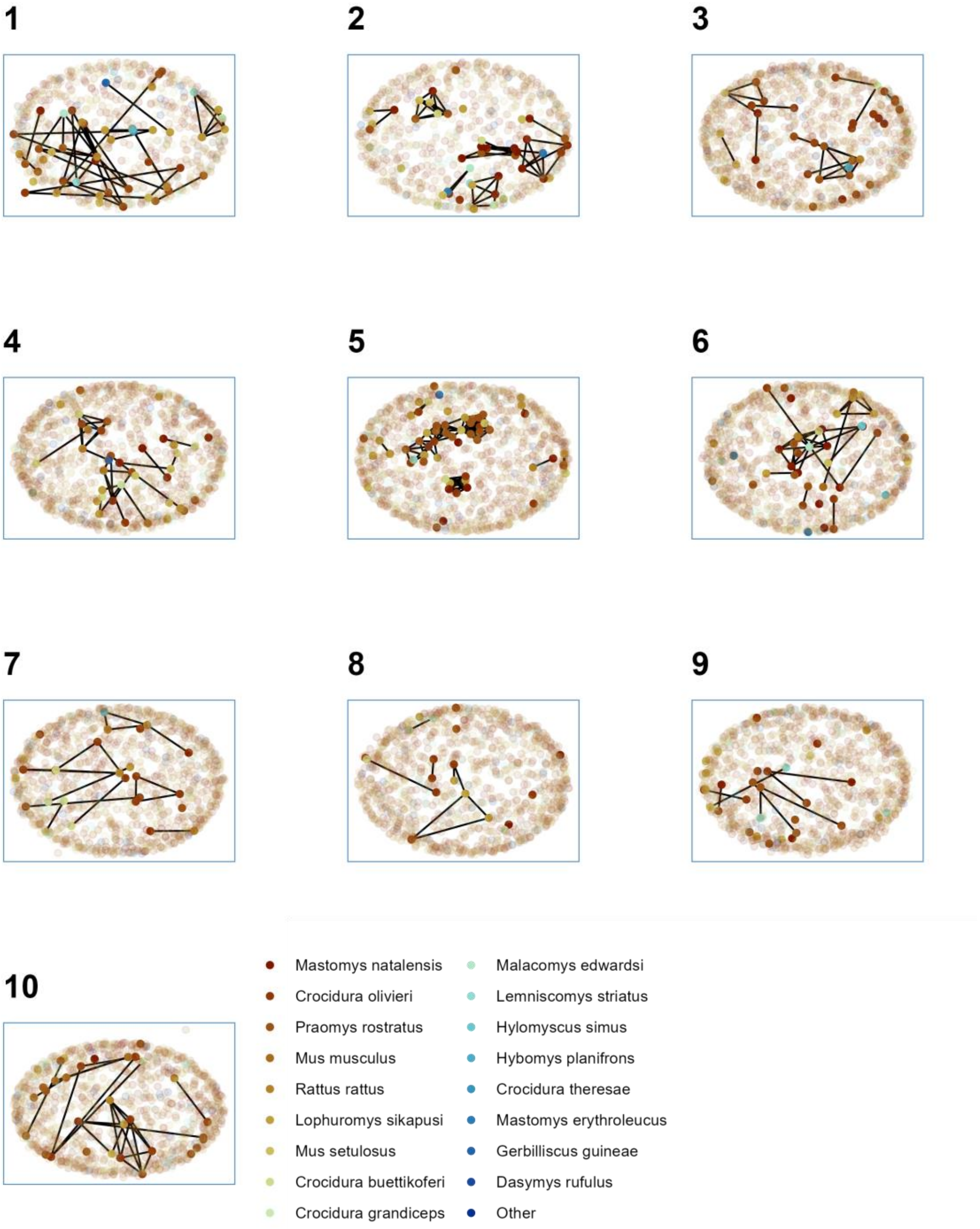

*Supplementary Figure 3.2: Networks produced from trapping sessions conducted in agricultural settings. Numbers refer to the trapping session. Observed individuals are shown in solid colours with colour referring to species. Unobserved individuals estimated from abundance modelling are shown in pale colours. Contacts between individuals (edges) are shown for observed individuals only.*

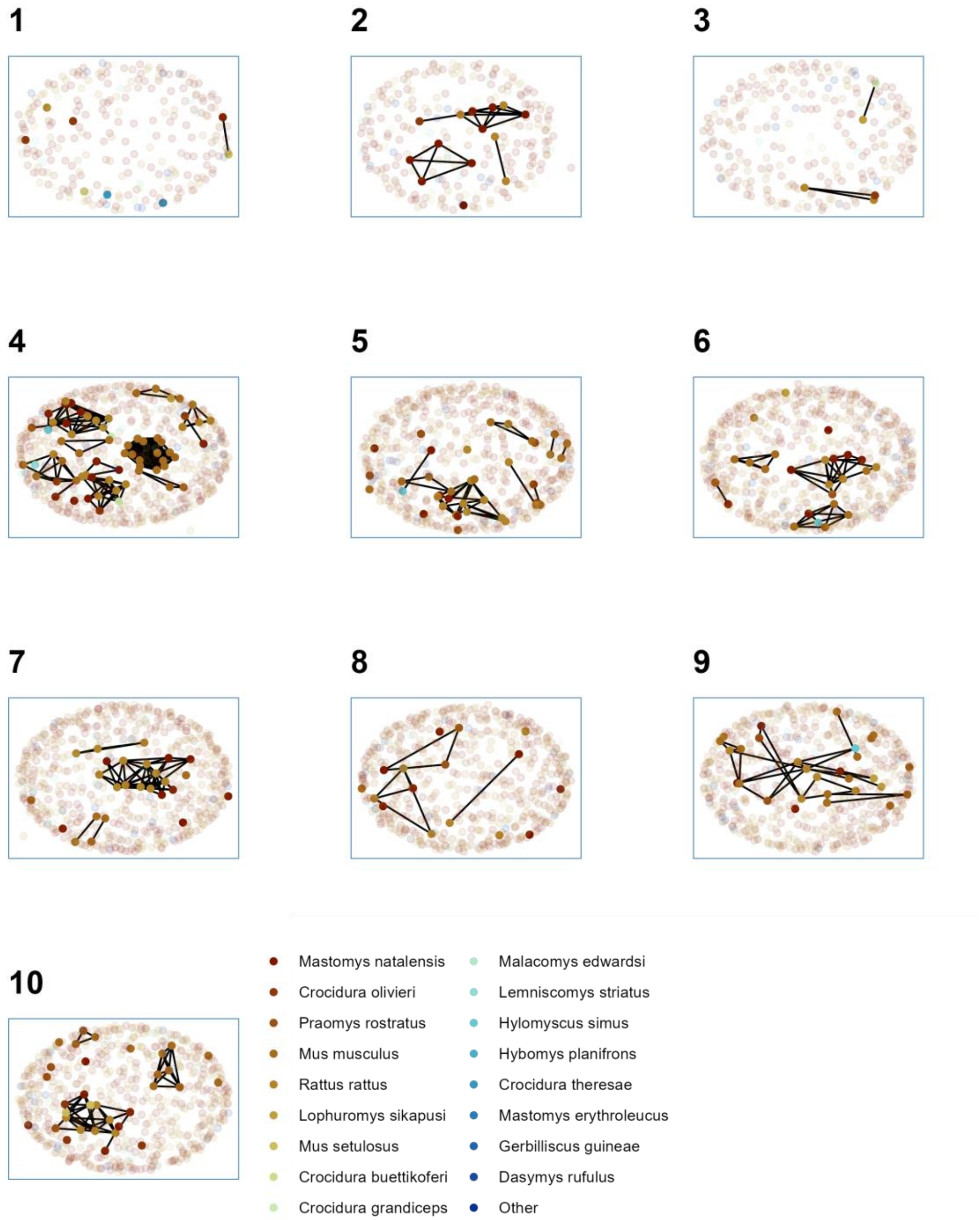

*Supplementary Figure 3.3: Networks produced from trapping sessions conducted in village settings. Numbers refer to the trapping session. Observed individuals are shown in solid colours with colour referring to species. Unobserved individuals estimated from abundance modelling are shown in pale colours. Contacts between individuals (edges) are shown for observed individuals only.*

### Supplementary Figure 4

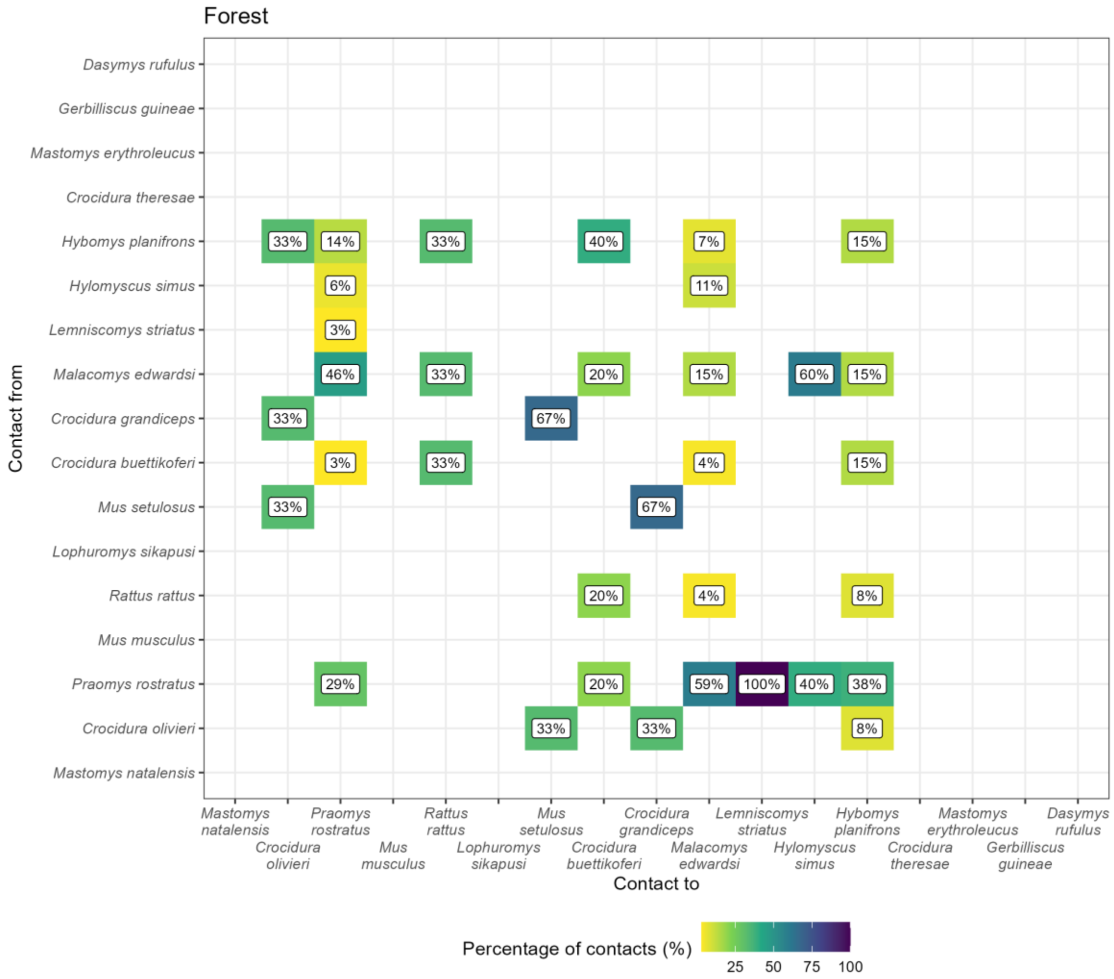

*Supplementary Figure 4.1: The proportion of contacts between individual small mammals in forest land use. Darker colours indicate increasing proportions of observed contacts to a species (Contact to) from named species (Contact from). Numbers in the cells correspond to the proportion of contacts to a species from a named species. For example, 56% of all contacts to Praomys rostratus are from other P. rostratus while 16% of contacts are from Malacomys edwardsi. Percentages sum to 100% in the* `Contact to` *axis, while they may exceed 100% in* `Contact from`*. Species are ordered by the total number detected in this study.*

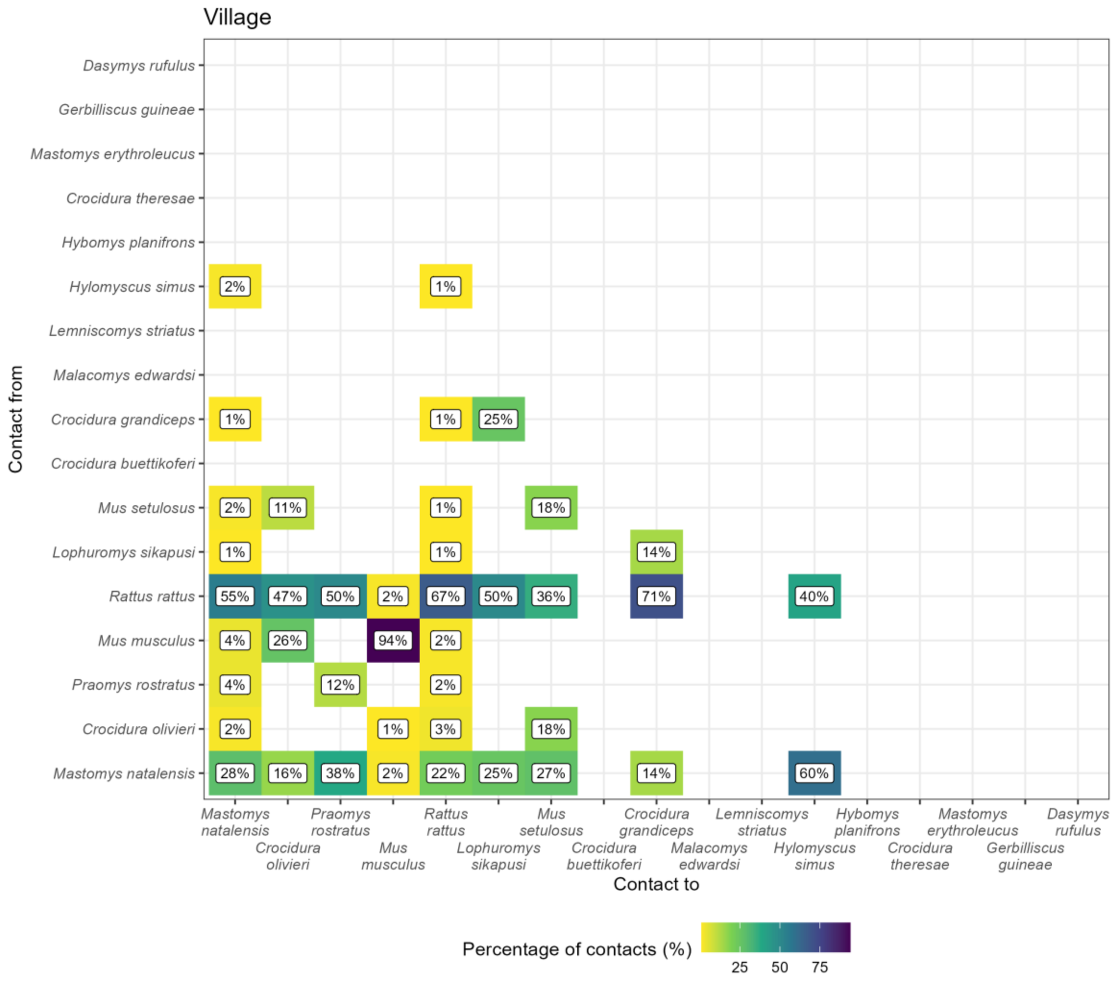

*Supplementary Figure 4.2: The proportion of contacts between individual small mammals in village land use. Darker colours indicate increasing proportions of observed contacts to a species (Contact to) from named species (Contact from). Numbers in the cells correspond to the proportion of contacts to a species from a named species. For example, 31% of all contacts to Mastomys natalensis are from other M. natalensis while 44% of contacts are from Rattus rattus. Percentages sum to 100% in the* `Contact to` *axis, while they may exceed 100% in* `Contact from`*. Species are ordered by the total number detected in this study.*

#### Supplementary Figure 5

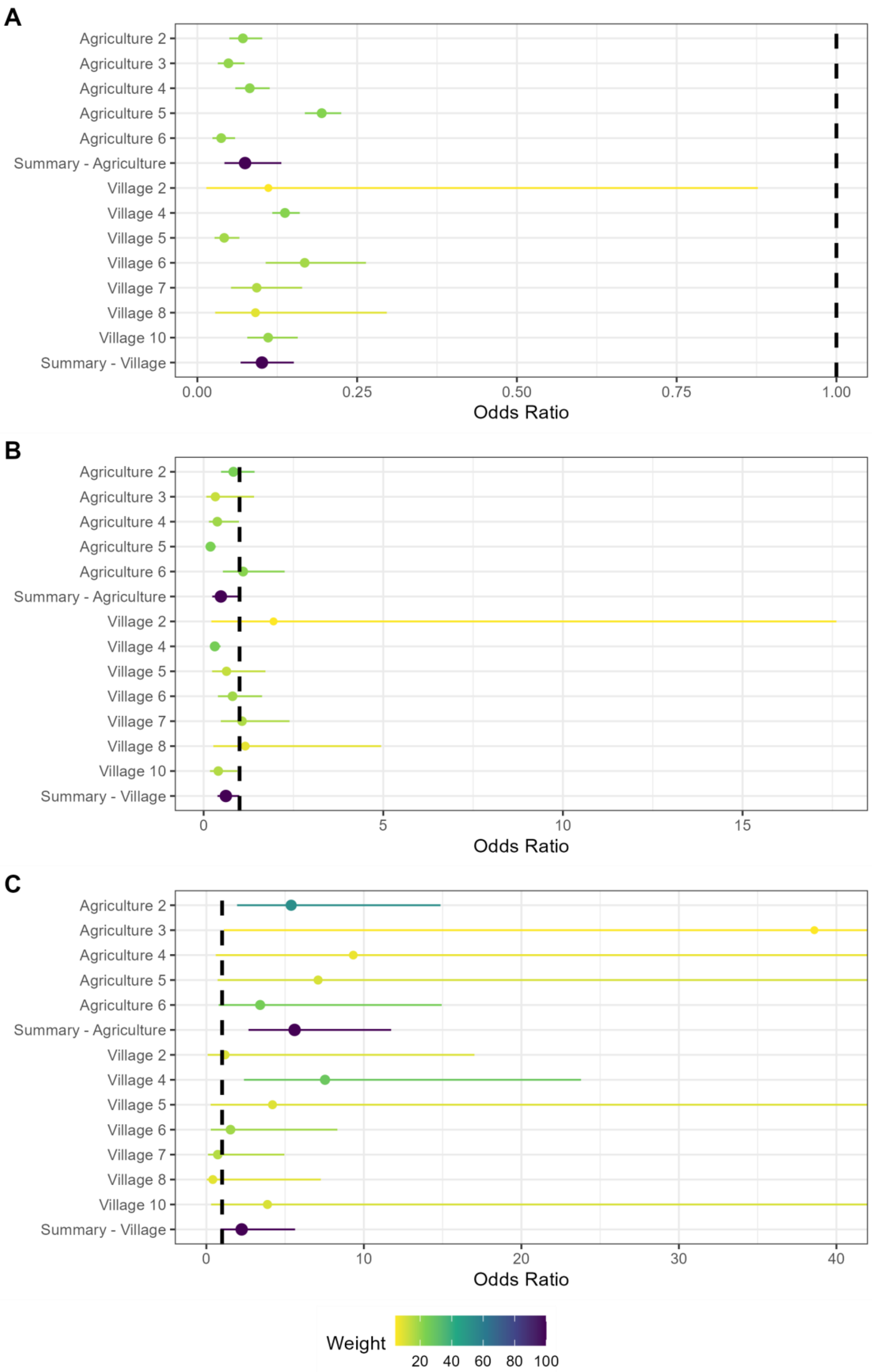
A
Agriculture 2
Agriculture 3
Agriculture 4
Agriculture 5
Agriculture 6
Summary - Agriculture
Village 2
Village 4
Village 5
Village 6
Village 7
Village 8
Village 10
Summary - Village
0.00
0.25
0.50
0.75
1.00
Odds Ratio
B
0
5
10
15
Odds Ratio
C
0
10
20
30
40
Odds Ratio
Weight
20
40
60
80
100

*Supplementary Figure 5.1: Random effects meta-analysis of ERGM network models reporting the odds of a contact being observed for M natalensis for the first sensitivity analysis by changing the buffer radius to 15m. A) The odds ratio of a contact being observed for M. natalensis in Agricultural or Village land use types. The finding of reduced odds of an observed edge is consistent with the main analysis. B) The odds ratio of a contact being observed between M. natalensis and an individual of a different rodent species. The finding of reduced odds of an interspecific edge is consistent with the main analysis. C) The odds ratio of a contact being observed between M. natalensis and another M. natalensis. This finding of increased odds of observing an intraspecific contact is consistent with the main analysis.*

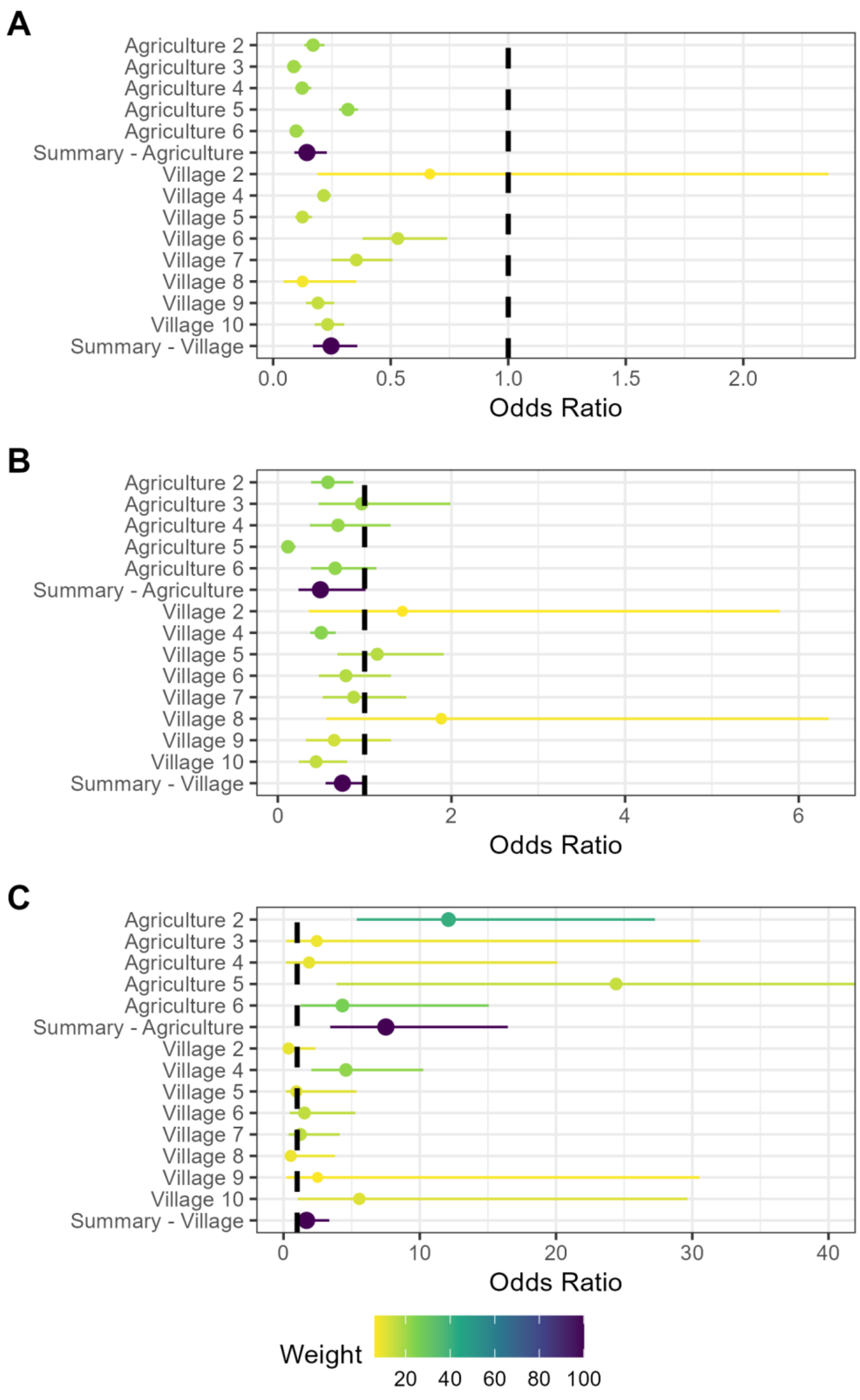

*Supplementary Figure 5.2: Random effects meta-analysis of ERGM network models reporting the odds of a contact being observed for M natalensis for the first sensitivity analysis by changing the buffer radius to 30m. A) The odds ratio of a contact being observed for M. natalensis in Agricultural or Village land use types. The finding of reduced odds of an observed*

*edge is consistent with the main analysis. B) The odds ratio of a contact being observed between M. natalensis and an individual of a different rodent species. The finding of reduced odds of an interspecific edge is consistent with the main analysis. C) The odds ratio of a contact being observed between M. natalensis and another M. natalensis. This finding of increased odds of observing an intraspecific contact is consistent with the main analysis.*

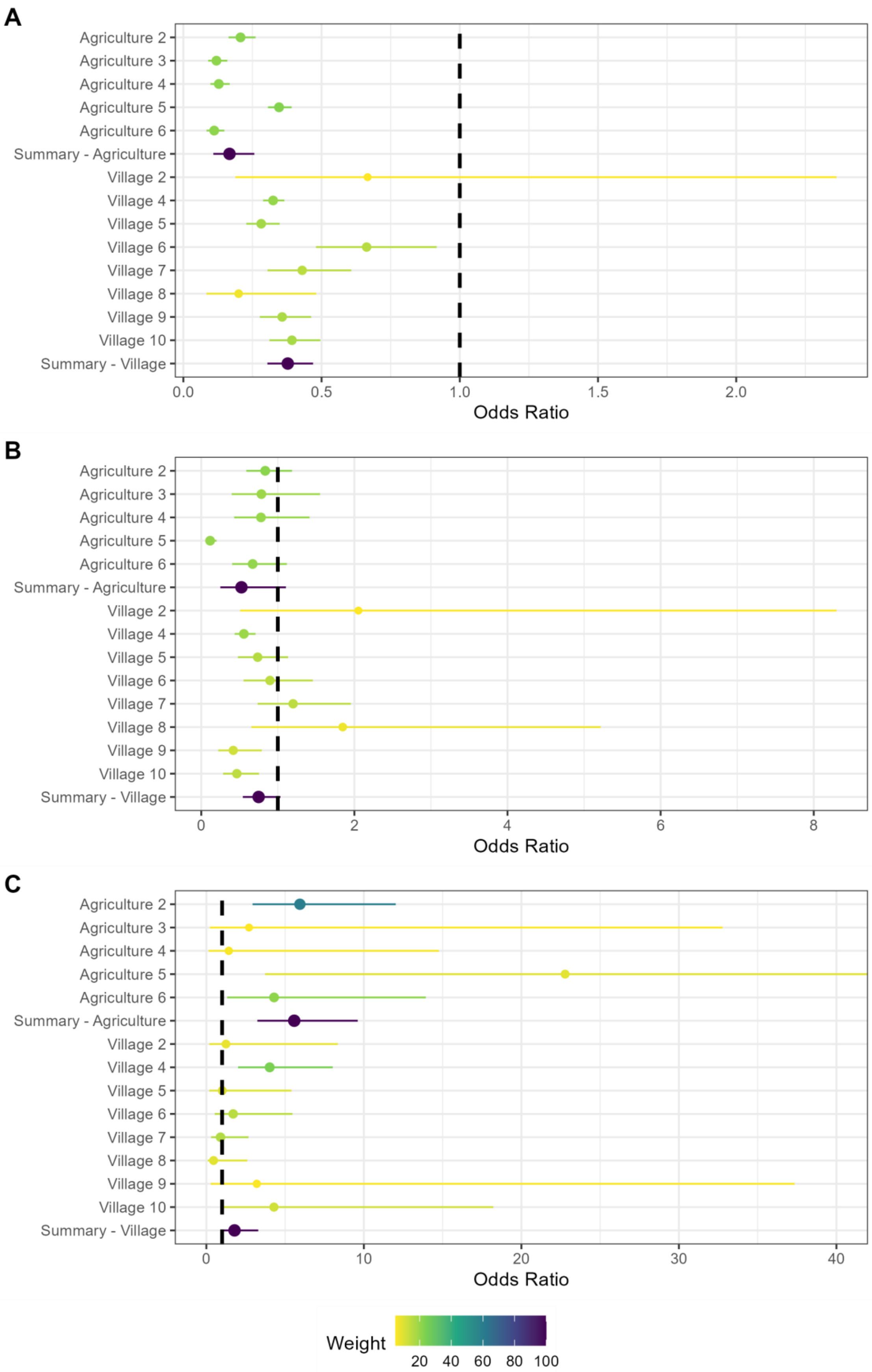
A
Agriculture 2
Agriculture 3
Agriculture 4
Agriculture 5
Agriculture 6
Summary - Agriculture
Village 2
Village 4
Village 5
Village 6
Village 7
Village 8
Village 9
Village 10
Summary - Village
0.0
0.5
1.0
1.5
2.0
Odds Ratio
B
Agriculture 2
Agriculture 3
Agriculture 4
Agriculture 5
Agriculture 6
Summary - Agriculture
Village 2
Village 4
Village 5
Village 6
Village 7
Village 8
Village 9
Village 10
Summary - Village
0
2
4
6
8
Odds Ratio
C
Agriculture 2
Agriculture 3
Agriculture 4
Agriculture 5
Agriculture 6
Summary - Agriculture
Village 2
Village 4
Village 5
Village 6
Village 7
Village 8
Village 9
Village 10
Summary - Village
0
10
20
30
40
Odds Ratio
Weight
20
40
60
80
100

*Supplementary Figure 5.3: Random effects meta-analysis of ERGM network models reporting the odds of a contact being observed for M natalensis for the second sensitivity analysis by changing the buffer radius to 50m. A) The odds ratio of a contact being observed for M. natalensis in Agricultural or Village land use types. The finding of reduced odds of an observed edge is consistent with the main analysis. B) The odds ratio of a contact being observed between M. natalensis and an individual of a different rodent species. The finding of reduced odds of an interspecific edge is consistent with the main analysis. C) The odds ratio of a contact being observed between M. natalensis and another M. natalensis. This finding of increased odds of observing an intraspecific contact is consistent with the main analysis.*
